# Adaptive Landscapes of *Plasmodium Falciparum* Dihydrofolate Reductase Reveal Pathways to Antifolate Resistance

**DOI:** 10.64898/2026.01.04.697492

**Authors:** Danny Muzata, Dwipanjan Sanyal, Deeptanshu Pandey, Soumyananda Chakraborti, Vladimir N. Uversky, Prajna Upadhyay, Sourav Chowdhury

## Abstract

Antifolate resistance in *Plasmodium falciparum* dihydrofolate reductase (pfDHFR) remains a major challenge for malaria control. To understand how this enzyme maintains function under antifolate selection, we developed PfPATH, a computational framework that integrates mutational fitness measurements, evolutionary interactions, and structural information to map adaptive trajectories. PfPATH adaptive walks reveal that evolution is constrained to a narrow ridge in sequence space defined by a small subset of residues. Most resistance mutations are individually deleterious and become viable only when supported by stabilizing changes, restricting adaptation to a few high-resistance pathways. Structure network analyses show that these mutations do not disrupt the global enzyme architecture but instead reorganize internal communication to preserve catalytic function. Consistent with this, molecular dynamics and free-energy analyses indicate that resistant variants remain stable while sampling a broader ensemble of low-energy conformational substates. Together, these results reveal a narrow and predictable route to antifolate escape.

## Introduction

Malaria remains one of the most challenging infectious diseases to control, partly because of the parasite’s ability to rapidly evolve resistance to frontline drugs (1). The mutation of *Plasmodium falciparum* dihydrofolate reductase (*Pf*DHFR) is one of the most challenging resistance mechanisms that minimize the efficacy of antifolate treatments (2). Antifolate drugs have long been used not only against malaria but also against bacterial infections and cancer to target enzymes in the folate biosynthesis pathway and block tetrahydrofolate production which is a necessary cofactor in nucleotide synthesis and cell growth (3, 4). In *P. falciparum*, dihydrofolate reductase plays a central role in folate metabolism by enabling the synthesis of DNA precursors essential for parasite proliferation and survival (5–7). This central role makes it an attractive target for antifolate drugs. This enzyme plays a vital process in the production of purines, thymidylate, and some amino acids. The effectiveness of antifolate medications, such as pyrimethamine and cycloguanil, basically hinges on this critical function (3). However, resistance is often acquired through point mutations in the *pfDHFR* gene that reduces drug binding without interfering with enzymatic function (5, 8). Mutations at residue positions 16, 50, 51, 59, 108, and 164 have been shown to reduce antifolate binding affinity (9, 10). Furthermore, combinations of these point mutations confer high levels of resistance. The frequently observed N51I/C59R/S108N triple-mutant haplotype results in substantial pyrimethamine resistance and is often accompanied by additional substitutions, such as I164L, giving rise to quadruple-mutant variants with even greater resistance (2, 11). Therefore, resistance in *Pf*DHFR does not arise solely from single substitutions, but from an evolved network of amino acid changes that collectively regulate protein stability, catalytic kinetics, and inhibitor binding (10, 12). These stepwise adaptations allow the parasite to navigate fitness trade-offs between maintaining function and evading drug pressure. Such complexities emphasize the remarkable adaptive plasticity of *P. falciparum*.

The evolutionary potential of *Pf*DHFR mutations highlights the threat and the ongoing risk of antifolate-based interventions in the treatment and prophylaxis of vulnerable populations (13, 14). While resistance in *P. falciparum* may involve mechanisms beyond mutation alone, understanding how mutational constraints preserve enzyme activity is a critical step toward understanding the resistance evolution. Although *Pf*DHFR-mediated resistance has been widely studied, the effects of resistance-associated mutations on enzyme structure, conformational dynamics and adaptive capacity remain poorly understood. A better way to frame this problem is to view the protein as a map of sequence space that encompasses all possible amino-acid variants. High-throughput mutational scanning and evolutionary studies have enabled the identification of both strongly constrained regions and more permissive, adaptable ones (15). This provides us with information on the evolutionary possibilities of the protein and its ability to respond to adaptive mutations. However, sequence-based approaches alone cannot fully explain the conformational plasticity and structural dynamics that are important for understanding adaptive protein fitness. Mutations can alter loop flexibility, active-site geometry, or long-range motions in ways that are not apparent from static sequence space analysis or crystallographic structures (16, 17). Hence, we combine evolutionary sequence analysis with all-atom molecular dynamics simulations to study proteins behavior at the atomic level. Molecular dynamics simulations provide a time-resolved perspective of the protein motions and reveal how enzymes search across protein conformations and how mutations influence those dynamics (18). In this study, we combine evolutionary sequence and structural features with all-atom molecular dynamics simulations of the apo wild-type *Pf*DHFR and known resistance mutants to construct a mechanistically informed fitness landscape. These complementary analyses form the basis of the *Pf*PATH framework, which links mutational fitness with conformational stability and dynamic constraints. Our approach provides unified evolutionary and biophysical perspective on antifolate resistance. We reveal how *Pf*DHFR adapts through constrained yet predictable mutational routes that may be interfered in the design of next-generation antifolate therapies.

## Results

### Evolutionary and Structural Features Inferred from Sequence Space

We investigated the sequence space of *Pf*DHFR to understand how changes in amino acids affect the protein’s fitness and stability. This revealed important aspects of the protein’s evolutionary history, showing that mutations at specific sites have different effects on stability, function and mutational tolerance. We also found the relationships between sequence variability and functional implications, especially at sites related to antifolate resistance. We first assessed evolutionary conservation by calculating Shannon entropy across all residue positions. With a *P. falciparum*-specific multiple sequence alignment, we identified that about 86% of the positions were conserved with most of the rest variable sites, particularly at the flexible loop regions (Fig. 1A). Among the most variable residue positions were N42, C50, S99, and I155 (Tables S1 and S2), all of which are associated with pyrimethamine resistance. To place these patterns in a broader evolutionary context, we also performed entropy analysis across diverse organisms. This comparison revealed six universally conserved positions (i.e. 30, 32, 38, 39, 91 and 104) each with zero entropy (Fig. 1A). This implies that residues at these positions are likely under strong evolutionary constraint. This suggests that substitutions at these positions could potentially disrupt structural integrity or catalytic function (Fig. 3B). Furthermore, most active-site residues exhibited low entropy suggesting limited tolerance to substitution and pointing to their importance for inhibitor binding and enzymatic activity.

**Fig. 1:**
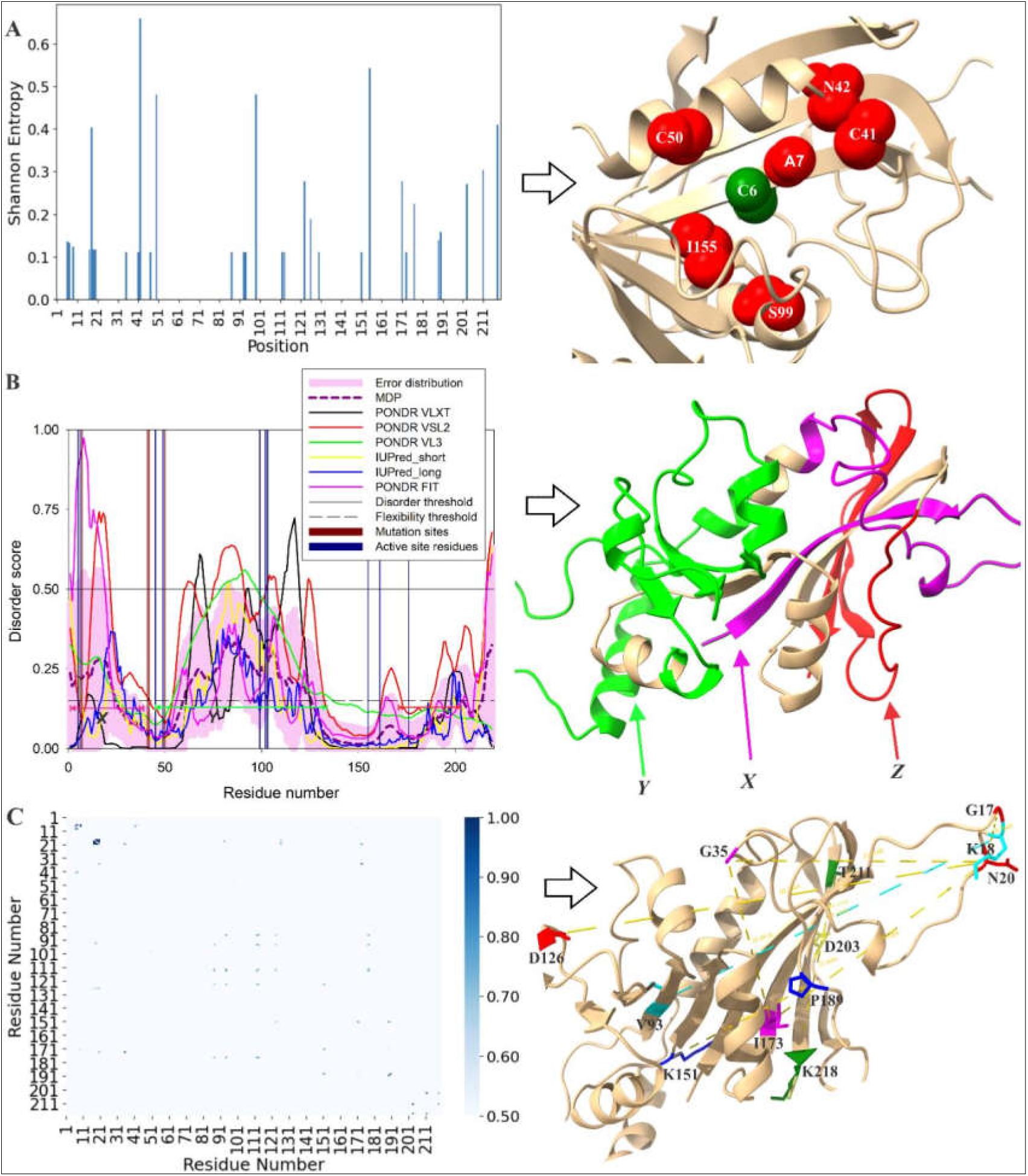
Sequence-derived features of *Pf*DHFR highlight regions of variability, intrinsic flexibility, and co-evolutionary coupling. **(A)** Shannon entropy values calculated from the *P. falciparum*–specific alignment highlights mostly conserved regions with a few variable positions. Highly variable residues occur in the loop regions (C41 and N42). Residue positions such as 6 (an active site position), 7 and 155 map to structured element like β-strands and while 50 and 99 map to α-helices. Thus, sequence variability is not limited to flexible loops. **(B)** Per-residue intrinsic disorder was assessed using six predictors: PONDR® FIT, PONDR® VSL2, PONDR®VL3, PONDR®VLXT, IUPred_short and IUPred_long. The mean disorder profile (MDP) captures their consensus. Residues with scores ≥0.5 were classified as disordered, whereas values between 0.15 and 0.5 indicate flexible but ordered regions. Although *Pf*DHFR is predominantly ordered, the insert highlights three segments that are relatively flexible labeled X (N-terminal region), Y (central region), and Z (C-terminal region) locally. **(C)** Mutual information (MI) analysis identified pairs of co-varying residues, with higher MI values (darker blue) indicating stronger evolutionary coupling. The insert maps represent high-MI pairs onto the 3D structure.

We then analyzed the flexibility of the *Pf*DHFR structure through six popular disorder predictors (20): PONDR® FIT (21), PONDR® VSL2 (22), PONDR® VL3 (22), PONDR® VLXT (23), IUPred Short (24), and IUPred Long (24) and generated its disorder profile. The result of these predictors was averaged to form a mean disorder profile that was used to give a consensus view of the protein’s disorder propensity. Intrinsic disorder usually is associated with a set of regions that can accommodate mutations as observed in the Shannon entropy profile. This adds to the evolutionary plasticity DHFR protein. As we found, this protein is largely ordered with most residues having disorder scores below 0.5, which points to the protein being stable. However, a few intrinsically disordered regions (≥ 0.5) were identified at positions 1–24, 60–125, 199–203, and 214–219 by the PONDR® FIT, PONDR® VSL2 and PONDR® VLXT predictors. The PONDR predictors (particularly PONDR® VSL2) are considered one of the most accurate predictors of disorder (22). Therefore, these disordered regions suggest points of potential mutational tolerance that can contribute to the protein’s adaptability (Fig. 1B). Although Shannon entropy described the general conservation trends, it failed to give us information on how individual residues may evolve in a coordinated manner. To analyze these inter-positional dependencies, we calculated Mutual Information (MI). Mutual Information measures the co-variation in pairs of residues and helps in determining where alterations could be compensatory to maintain structure or functional integrity. The MI heatmap (in Fig. 1C) indicated that there were several strong co-evolutionary signals, denoted by the dark blue spots, at the N-terminal, central region and the C-terminal end. Some of the high-MI pairs were mapped onto the 3D structure which illustrate both short-range contacts and long-range dependencies spanning distant structural elements. These strongly co-varying residues later appear interacting with critical functional regions (like mutation sites and active-site residues) as shown in the MI networks in Fig. 2.

**Fig. 2.**
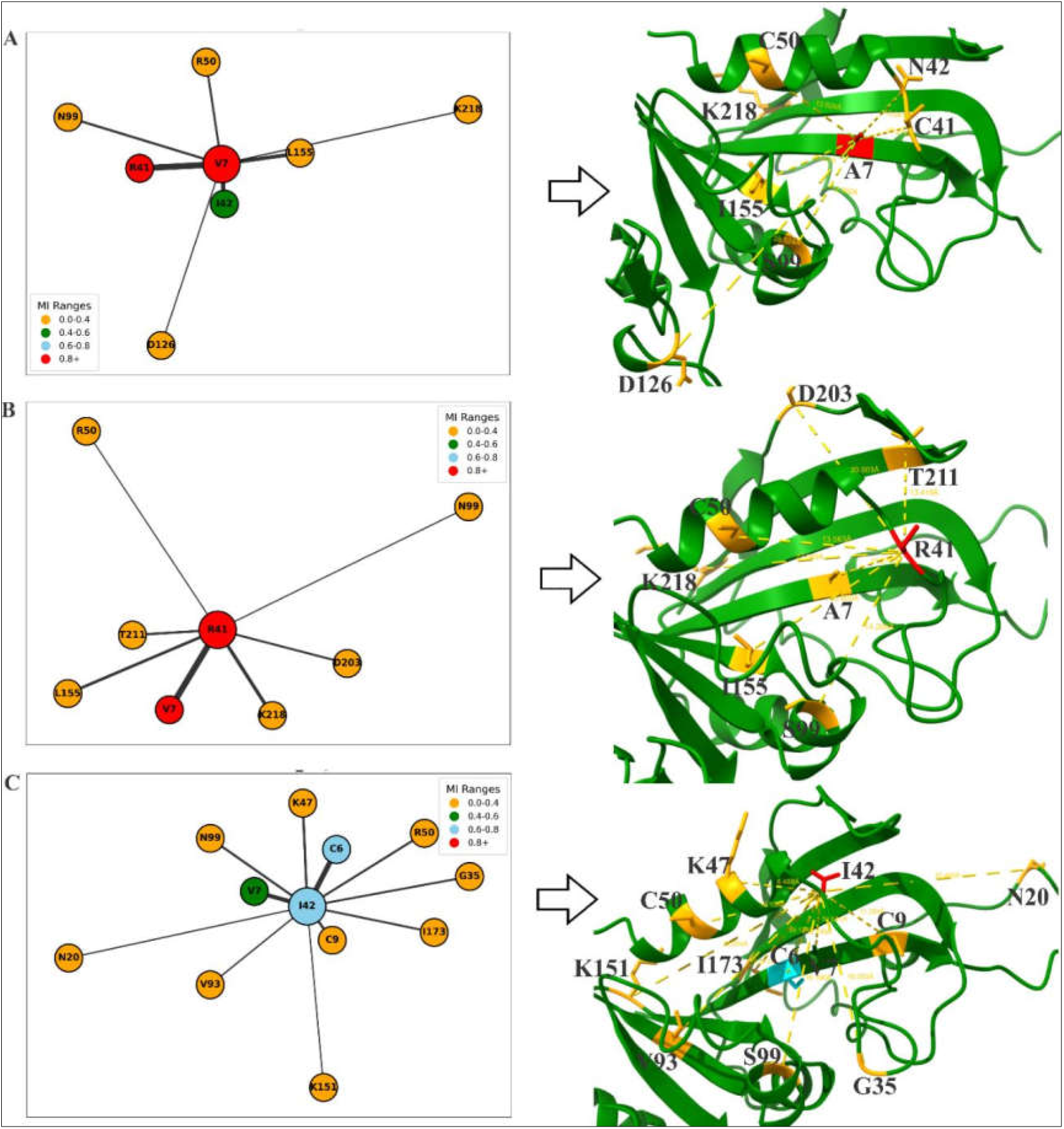
Local Mutual Information networks centered on key *Pf*DHFR mutation sites reveal extensive co-variation across distant residues. **(A)** The network around mutation site A7 shows strong co-variation with multiple residues. The node colors reflect MI strength and the inserts on the right illustrate how these residues are distributed across β-strands, helices, and loop segments. **(B)** The R41 network revealed a wide interaction spread extending across the protein fold. Several co-varying partners overlap with the A7 network. **(C)** Mutation site N42 also exhibits moderate co-variation with residues distributed across the fold, including positions that lie close to the catalytic core. The insert shows these connections bridging distant regions of the protein.

With the correlations seen in the MI heatmap, we further examined the local interaction networks formed around these individual mutational sites (Fig. 2). The networks revealed more pronounced and extended patterns of co-variation of these sites. They create an interaction hub with distributed residues across different regions in the protein structure. The respective 3D inserts also show that these interactions span both short-range and long-range distances. Hence, co-variation is not driven by spatial proximity. For example, the A7 interaction network (Fig. 2A) revealed co-varying partners, many of which sit in distant regions in the protein. It had the strongest correlation with residue C41, a position on which three resistance substitutions by Arginine, Tyrosine and Asparagine have been reported (8, 10). A similar interaction network was observed for R41 displaying wide-reaching interactions across both β-strand and loop regions. N42 and C50 (Fig. S4) even formed co-variation links with residues in the central catalytic core. In contrast, position I155 showed weaker but still notable interactions, suggesting a more modest degree of evolutionary coupling (Fig. S4). Table S3 summarizes the MI values and Cα–Cα distances for each network. Across the networks, mutation sites such as N42, C50 and I155 were linked to other mutation-associated positions and residues situated near the active-site region. This pattern suggests that resistance evolution in *Pf*DHFR may be partly coordinated across a distributed set of these interacting residues. The distances between co-varying residue pairs covered a wide range. This indicates that strong MI correlation does not depend on how close the residues are in 3D space, and that some residues can co-evolve via long-range evolutionary coupling. Such residues could serve as intermediate or supporting positions in compensatory pathways.

### Network-based analyses reveal compensatory hubs and shared structural modules linking active and mutation-prone residues in *Pf*DHFR

To understand how the co-variation patterns identified in the MI map are organized within the protein structure network, we analyzed the residue–residue correlation network using maximal clique analysis (25, 26) and community detection (27). These approaches describe the organization of the protein at two levels. Maximal cliques capture very tightly coupled micro-modules where all residues are connected to each other. The community detection captures larger dynamic regions that move together on average. These methods highlight which residues form local compensatory clusters and which belong to broader functional modules. In the maximal clique analysis, we identified 270 cliques across the *Pf*DHFR network. Several mutations and active-site residues were consistently located in the same dense cliques. Residues 41, 42, 50, 99 and 155, as well as active-site positions 5, 6, 161, 176, 102 and 103, belonged to the same large 8-member clique. The summary Tables S4 to S9 shows that positions 7, 99 and 155 appeared across dozens of cliques. These residues co-occur with active-site positions and with other mutation sites suggesting that they may help buffer the structural consequences of resistance-associated substitutions. For example, residues 42 and 50 co-occurred with active-site residues 45 to 49 in several cliques. These patterns support the interpretation that *Pf*DHFR leverages locally connected clusters of residues to maintain catalytic geometry while tolerating destabilizing changes. Community detection further divided the residue network into nine larger sub-structures. Mutation and active-site residues again appeared together in several communities. Residues 41, 42 and 50 are grouped with active-site residues 45, 46, 48 and 49 in Community 5 while residues 99, 102 and 103 clustered in Community 1. These communities represent broader segments of the protein that exhibit coordinated motion. All these analyses show that *Pf*DHFR resistance evolution is supported by both fine-scale and larger-scale structural coordination.

### Deep mutational scanning revealed that *Pf*DHFR is globally constrained yet locally flexible, with conserved core and catalytic residues displaying strong intolerance to substitutions

To quantify the functional consequences of amino acid changes across the sequence, we carried out a Deep Mutational Scanning (DMS) (28). This technique measures the fitness impact of every possible single-residue substitution. This allows us to map the mutational landscape of the protein and distinguish regions that are highly constrained from those that tolerate changes. In the heatmap (Fig. 3C), beneficial substitutions are shown in shades of blue, whereas deleterious ones appear in red. The results show that different positions vary widely in how much mutation they can tolerate. At residue position D1 for example, most substitutions are beneficial. This means substitutions at this position can accommodate the changes without compromising function. However, substitutions at the neighboring positions were predominantly deleterious. Distinct regions of mutational intolerance were also observed across residues 30–53 and 90–196, whereas residues 54–90 (a loop–helix segment) exhibited comparatively higher mutational permissiveness. Two positions, 156 and 157, showed extreme intolerance. Nearly all substitutions at these sites were deleterious and this was consistent with our Shannon entropy analysis which identified them as highly conserved sites. Active-site and resistance-associated positions also followed expected patterns. Residue 7 was strongly deleterious across most substitutions, while residues 41 and 42 exhibited beneficial profiles. Conversely, residue 99 and residue 155 were strongly deleterious for most substitutions. The apparent contradiction between the fitness costs at these positions and natural occurrence of resistance mutations at the same sites could be explained by the compensatory or buffering mechanisms as observed in our MI network analysis (29).

**Fig. 3:**
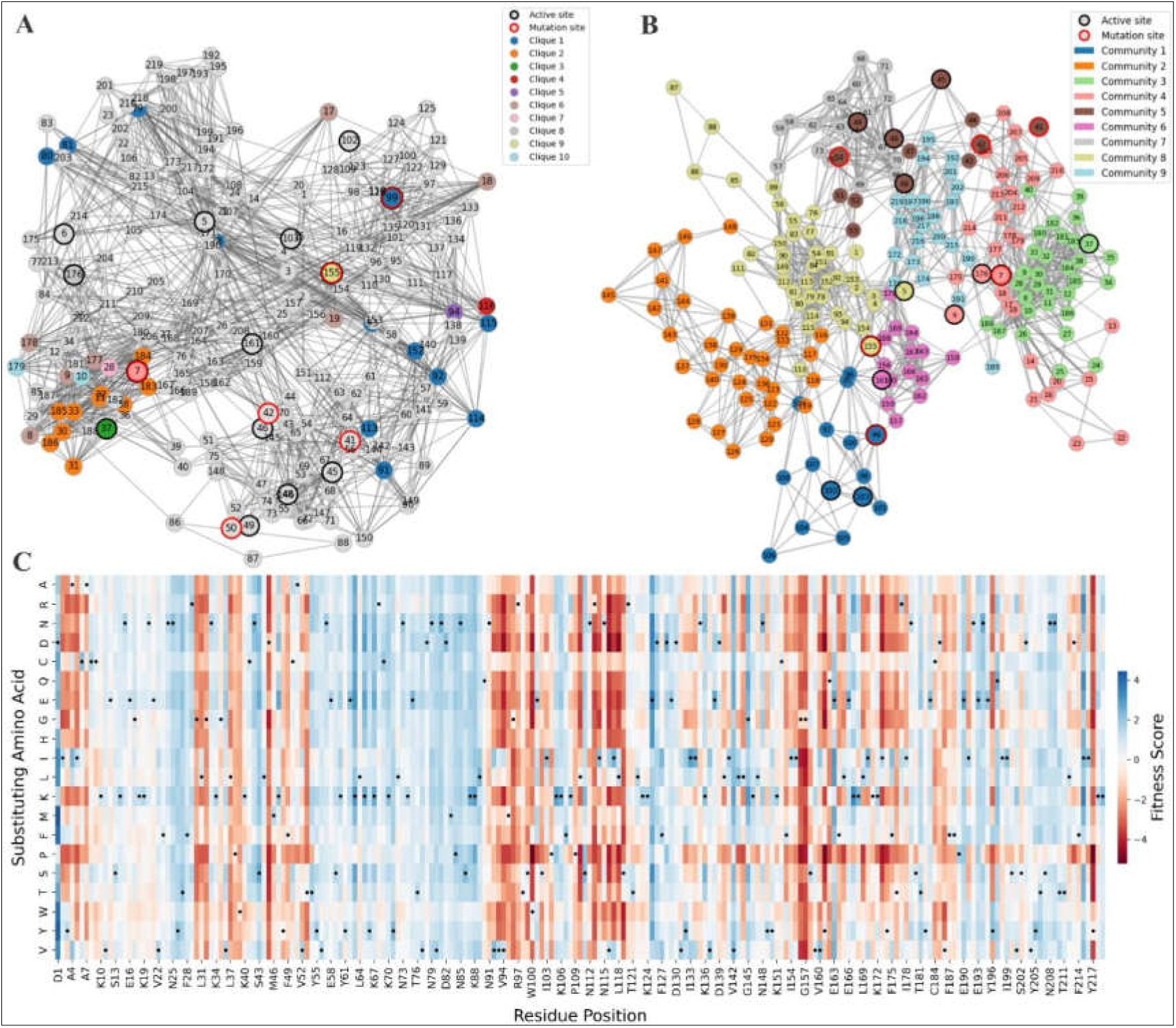
Network topology and mutational scanning highlight the structural basis of *Pf*DHFR adaptability. **(A)** Maximal clique analysis shows the top ten color-coded cliques in the residue–interaction network. Active-site residues (black outlines) and mutation sites (red outlines) frequently appear within the same cliques. **(B)** Shows community detection of the DHFR interaction network. Nodes represent residues and edges represent correlated motions. Nine communities were identified. Residues that function as both catalytic and mutation hotspots are marked with both outlines and can be seen belonging to the same community in some cases. **(C)** Deep mutational scanning heatmap displays the predicted fitness effects for all single amino-acid substitutions across the protein fold. Blue shaded regions indicate tolerated or beneficial substitutions, whereas red shades indicate deleterious effects.

### Structural Features Inferred from 3D Protein Structure

Our sequence-based analyses identified differences in residues that tend to change together. These patterns, as informative as they are, do not indicate how the protein copes with these substitutions within the three-dimensional structure. Specifically, pairwise evolutionary signals are unable to tell the difference between the change which was caused by local packing and that which was caused by long-range dynamic communication. To address this, we looked at the correlated motions and structuring of dynamic sub-networks based on the dynamic cross-correlation matrix (DCCM) (30). The WT contact map (Fig. 4C) exhibited a landscape with no large positive correlated regions but had a predominantly weakly correlated. These were mostly localized in the N-terminal and the C-terminal tail. Eight communities were detected and as we observed in our previous analyses, many active-site residues and mutation-prone residues belonged to the same communities and not isolated. This demonstrates that the important catalytic and mutation-sensitive residues are involved in a distributed communication scaffold. We also noted that all mutants modified the WT network differently. The C41R mutant weakened some of the contacts identified and resulted in more anticorrelated patches throughout the DHFR fold (Fig. S6). Because of this mutation, the network reorganized itself into ten communities where C41 clustered together with the active size residue C50 in community 5. The other active-site residues moved into different communities compared to what we observed in the WT. On the contrary, the N42I mutant was seen to strengthen some contacts especially in the N-terminal. The network expanded into eleven communities and the mutant remained within the 41–55 structure block with active-site residues 45–49 (Fig. S6). C50R and S99N mutations resulted in even more anticorrelations. C50R also existed in the same united structure block with N42 and C41. Although long-range couplings were weakened, new short-range contacts were obtained especially between the stretch of residue position 75 and 85. S99N clustered with the active sites 102 and 103 although they moved into a smaller community. A lot of correlations with the C-terminal were lost. The I155L mutant showed the appearance of strong positive correlations around residues 68–79 as well as prominent anticorrelation patches. It is grouped with residues sites 99, 102 and 103. A residue-level view of how these local changes reorganize dynamic communities across the trajectory is shown in Fig. S7. Mutations in all systems neither isolated residues nor broke down the network. Rather, they reshaped the communication pathways in a distinct manner. The WT enzyme has a unified, hub-based structure that has established communication channels. Mutations either destroy or form new short or long-range stabilizing interactions. The result is a list of other communication solutions that enable the protein to trade-off between catalytic demands and conformational changes related to resistance.

**Fig. 4.**
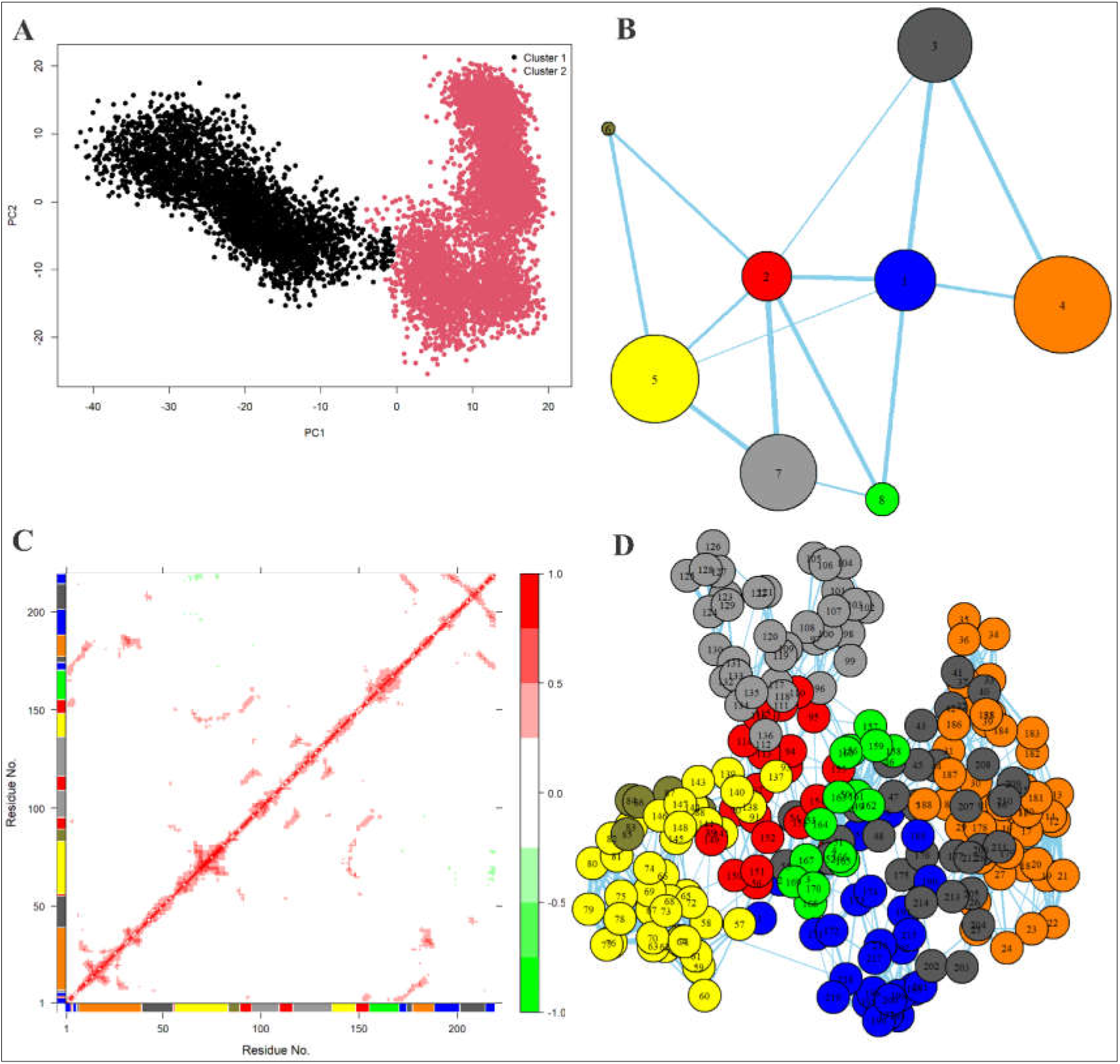
Local flexibility and correlated motions in WT *Pf*DHFR. **(A)** Principal component analysis shows that the WT enzyme samples two dominant conformational clusters indicating transitions between long-lived substates. **(B)** Community network representation of the WT dynamic correlation map. Each node represents a community of residues that move together, with node size proportional to community size and edge thickness indicating the strength of inter-community coupling. **(C)** The contact map of WT shows positive correlations (red) and negative correlations (green) which reveal patterns of cooperative and opposing motions across the fold. The color bars along the axes indicate community assignments for each residue. **(D)** All-residue community network in which each node corresponds to a single residue and is colored according to its community membership.

### Molecular Dynamics Reveal a Stable Wild Type Fold and Mutations Increase Local Flexibility

Molecular dynamics (MD) simulations of the apo form of WT *Pf*DHFR were performed to complement the sequence-based analyses and to understand how the protein behaves in 3D over time. Although entropy and co-variation describe mutational tolerance and coupling, molecular dynamics help show how the protein fold is able to accommodate these sequence-level changes by intrinsic motions. Across the one-microsecond trajectory, the WT enzyme remained globally stable. After initial equilibration (about 100–350ns), the structure settled into a stable conformation by roughly 550ns and maintained this state for the rest of the run. The RMSD plateaued around 4Å with only a single transient peak near 5Å. This indicated no unfolding or large-scale destabilization (Fig. 5A). The radius of gyration (Fig. S9A) dropped slightly from approximately 1.98 to 1.85nm suggesting a slight compaction and the number of intramolecular hydrogen bonds remained steady at roughly 145–150 throughout the simulation (Fig. 5C). Per-residue fluctuations showed a clear division between a rigid catalytic core and highly dynamic loop regions. Loop 1 (residues ∼20–35), Loop 2 (∼85–100) and the C-terminal tail (∼180–200) (Fig. 5B) displayed significant motions in the mutants while the catalytic core remained tightly constrained. Solvent Accessible Surface Area (SASA) was consistent with the small compaction depicted in the radius of gyration (Fig. 5D). Similarly, in all the resistance mutations, the global fold of *Pf*DHFR remained stable. But the local disruptions in the structure, particularly in the flexible loops and the adjoining segments were introduced by these mutations. In the WT simulation, loop motions reached peaks of around 6Å, whereas all mutants showed stronger fluctuations, often exceeding 7–8Å. The N42I mutant produced the strongest increase in flexibility, especially across Loop 2, where it displayed the highest RMSF values of all variants. This means the mutation can potentially cause destabilization of adjacent segments but the protein balances it by redistributing the movement on the surrounding loops (10). I155L showed a similar pattern. The core itself was stable, yet surface-exposed loops were more mobile and the radius of gyration (Fig. S10) was a little bit higher which suits its purpose of being a permissible mutation in multi-mutation resistance backgrounds (10). S99N exhibited a low degree of local flexibility in the vicinity of the binding pocket that is in line with previous results that have associated this mutation with pocket adjustments and a modified interaction with ligands (31). The other resistance mutations also enhanced local flexibility about their respective substitution sites without adversely affecting the structural integrity. Despite the specific patterns of fluctuations being varied in them, they all had increased loop mobility compared to the WT with no apparent destabilization. These findings demonstrate that the structure of *Pf*DHFR is robust, and mutations that confer resistance always reshape local dynamics. They do not break the fold but instead destabilize their immediate neighborhoods and redistribute flexibility across surface loops that surround or communicate with the catalytic region. This observation is consistent with our co-variation analyses which showed that mutation sites and active-site residues form compensatory hubs.

**Fig. 5.**
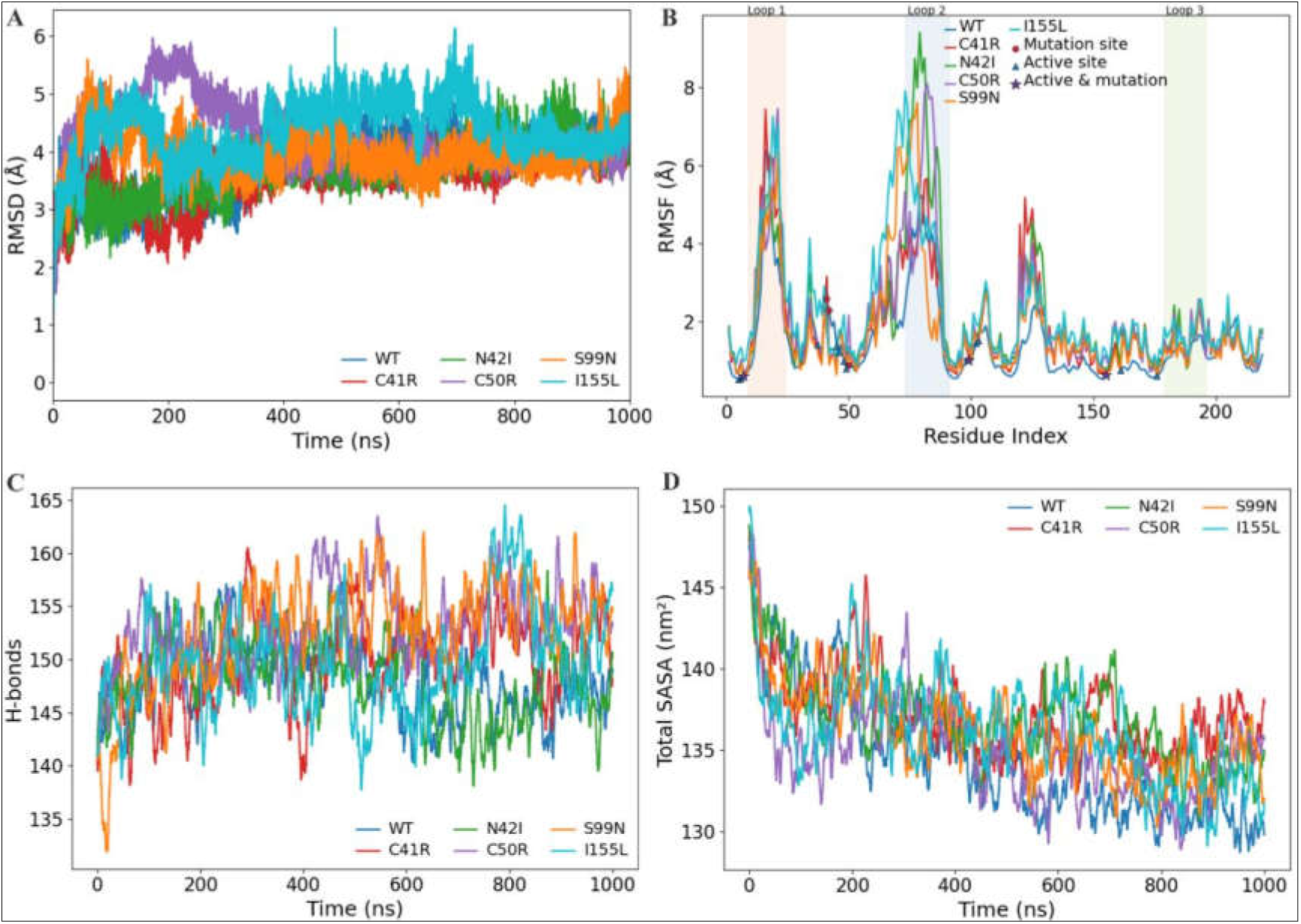
MD comparison of WT and mutant *Pf*DHFR shows stable global folds with mutation-specific increases in loop flexibility. **(A)** All variants equilibrated within the first ∼350 ns and remain folded throughout the simulation. RMSD differences reflect conformational shifts rather than unfolding. **(B)** RMSF profiles showed that the same three loops are flexible in all systems, but the mutants consistently display higher fluctuations than WT. **(C)** Hydrogen-bond counts remain within a similar range for all variants indicating preserved overall packing. **(D)** Total SASA stabilizes between ∼130–140 nm² across systems with only minor differences in compactness.

### Free-energy landscapes reveal a rugged WT funnel and flattened, multi-basin mutant ensembles

The MD simulations demonstrated that *Pf*DHFR has a stable core with very flexible loop regions. However, these motions could only describe the movement of the protein, but not the energy cost and occupancy of each state. To capture this thermodynamic dimension, we constructed free-energy landscapes (FELs) by projecting all trajectories onto the same principal component plane so that the same collective motions apply to both the WT and all the variants. Free energy was calculated from the probability distribution using ΔG = –kBT ln p, where kBT is the thermal energy (10). The 3D surfaces (Fig. 6) and corresponding 2D contours (Fig. S11) map how often each conformation is visited. The valleys represent low-energy, frequently sampled states, while elevated regions correspond to high-energy conformations rarely accessed. The WT enzyme forms a rugged but focused funnel with most sampling concentrated around a single dominant low-energy basin. About 35% of frames fall within 2 kJ/mol of the minimum, 94% within 5 kJ/mol and nearly all within 10 kJ/mol, which indicates a well compacted native ensemble. All the five mutants span a similar overall energetic range (≈31–32 kJ/mol). However, they differ markedly in the shape of their low-energy regions. Each mutant shows a flatter minimum which indicates a redistribution of conformational probability into broader, shallower basins rather than an expansion of the total energy range. This effect was most pronounced in S99N, with approximately 71% of frames lying within 2 kJ/mol of the minimum, followed by C50R (∼67%) and N42I (∼61%). I155L showed intermediate broadening, while C41R remained most similar to the WT. Such differences show that resistance-associated mutations restructure the organization of accessible conformations without altering global stability. The wider and less deep basins reduce the energetic barriers between the nearby substates. This enables the mutants to be able to access a broader range of low-energy conformations. This enhanced accessibility offers a thermodynamic pathway through which *Pf*DHFR can retain catalytic activity but be less vulnerable to antifolate inhibition.

**Fig. 6:**
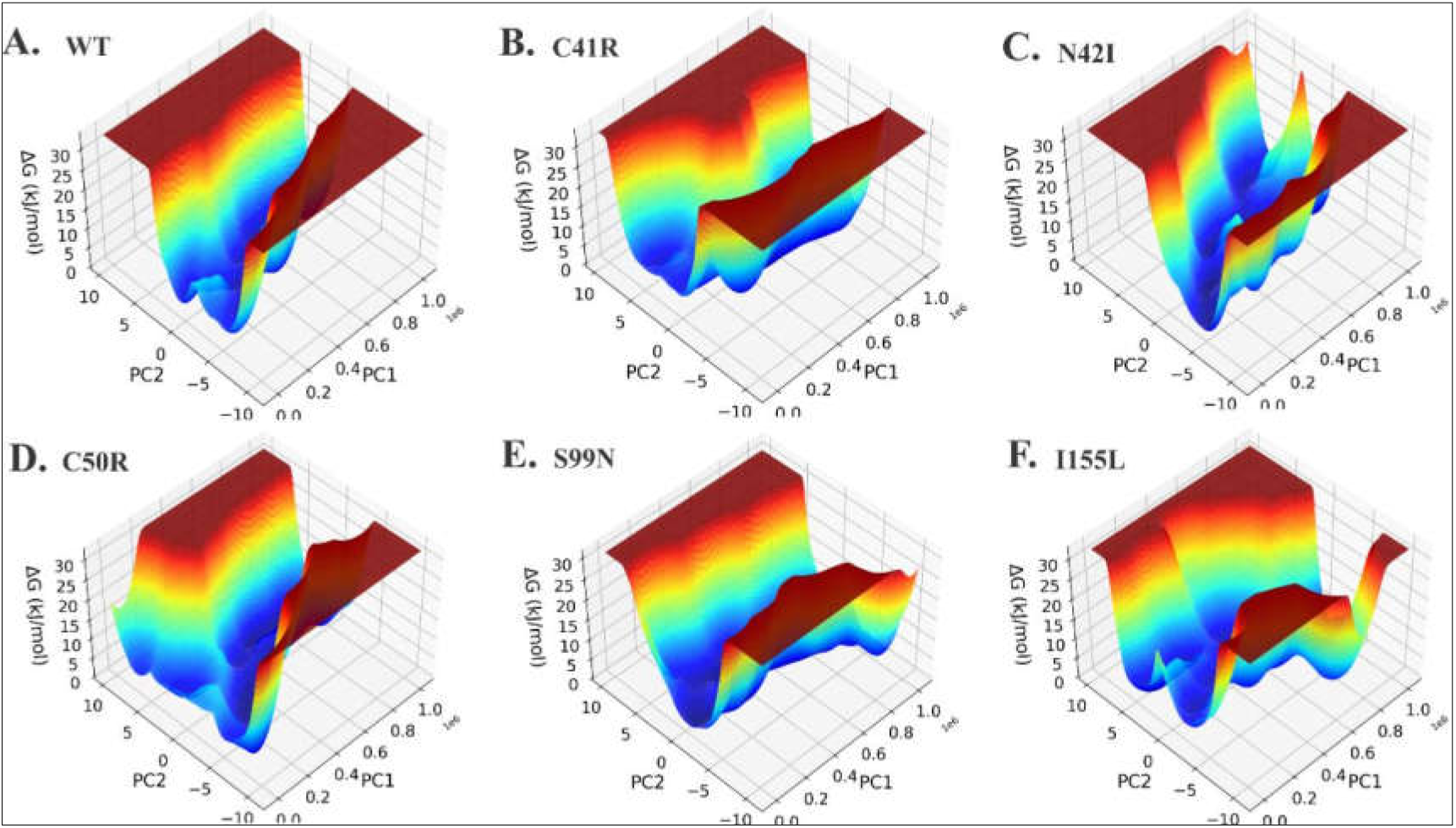
Free-energy landscapes of *Pf*DHFR reveal that mutations flatten the WT funnel, leading to more flexible and adaptive conformations. **(A)** WT shows a steep, well-defined funnel with deep minima which is characteristic of a rigid folded ensemble. **(B)** C41R displayed a broader, saddle-like surface with multiple shallow dimples **(C)** N42I presented an uneven landscape with connected minima, indicating increased substate diversity. **(D)** C50R formed a flattened, near-degenerate basin allowing easy movement between similar conformations **(E)** S99N had the flattest surface with multiple equivalent minima linked to loop flexibility. **(F)** I155L showed a wide main basin with shallow side basins separated by low ridges. The shifts between nearby conformations are easier than in WT.

### Adaptive-walk simulations reveal a latent compensatory ridge of *Pf*DHFR adaptability

To understand how *Pf*DHFR navigates the rugged fitness landscape, we developed the *Plasmodium* Fitness Pathway Analysis of Trajectories in Heterogeneous Landscapes (*Pf*PATH) model. This model simulates stochastic adaptive walks through a composite fitness landscape, which combines evolutionary sequence constraints, co-evolutionary coupling, protein stability, and experimentally derived biochemical and kinetic measurements. These adaptive walks induce an initial mutation at the starting position and the algorithm then explores more fit neighbors by using the composite fitness function. Each bundle initially explored distinct mutational solutions depending on the first-step substitution. Every climb quickly narrowed down to a common high-fitness plateau, which indicates a segment of sequence space that is very accessible and resilient to the selection process. Importantly, many of the same compensatory substitutions recurred across the independent walks and across different starting residues. This repeated convergence highlights what we term as “latent compensatory ridge”, a narrow set of positions that repeatedly serve as stabilizing, path-enabling intermediates regardless of where adaptation begins. Table 1 below shows this set of recurring residues. Positions such as V36, R29, K30, L31, C84, F187, R177, P38, L212, N209, T54 and neighboring residues repeatedly appeared in high-fitness trajectories. These sites occupy dynamic nodes that enhance stability, flexibility or co-operative behavior. Their repeated emergence implies that *Pf*DHFR relies on these locations as buffering points that allow costly resistance mutations to be accommodated without collapsing catalytic function.

**Table 1.** Recurrent stabilizing residues supporting adaptive pathways in *Pf*DHFR.

| Rank | Position | Evidence in Adaptive Walks |
| --- | --- | --- |
| 1 | I215 | Very frequent |
| 2 | F214 | Frequent |
| 3 | I216 | Frequent |
| 4 | P189 | Frequent |
| 5 | R177 | Regular |
| 6 | K123 | Regular |
| 7 | T211 | Occasional |
| 8 | V94 | Occasional |
| 9 | L31 | Occasional |
| 10 | R29 | Occasional |
| 11 | V36 | Rare but recurrent |
| 12 | P38 | Rare but recurrent |
| 13 | G30 | Rare but recurrent |
| 14 | C184 | Rare but recurrent |
| 15 | F197 | Rare but recurrent |
| 16 | L212 | Rare but recurrent |
| 17 | N209 | Rare |
| 18 | T54 | Rare |

We also examined how these resistance-associated positions are selected and maintained during long-term evolution under fluctuating drug pressure. Simulations using a Wright–Fisher evolutionary model (32) were performed with a population size *N* of 10,000 per-site mutation rate (µ = 10^-5^) and a simple drug-cycling regime consisting of three generations with pyrimethamine “on” followed by one generation “off” repeated throughout the run. For each generation we tracked the fraction of the population carrying a non-wild-type amino acid at six key positions, that is, 7, 41, 42, 50, 99 and 155 which correspond reported resistance–associated sites (Fig. 7F). The trajectories revealed a clear temporal hierarchy of mutations at known resistance sites. The earliest and strongest sweep occurred at position 42, which climbed from near zero to >0.7 frequency by generation ∼8,000 and eventually approached fixation. Position 41 rose later and more modestly, forming a broad transient peak centered around generations 10,000–11,000. Positions 50 and 155 showed only weak, delayed increases at low frequency near the end of the simulation. In contrast, positions 7 and 99 remained essentially neutral with mutation frequencies fluctuating around zero across generations.

**Fig. 7.**
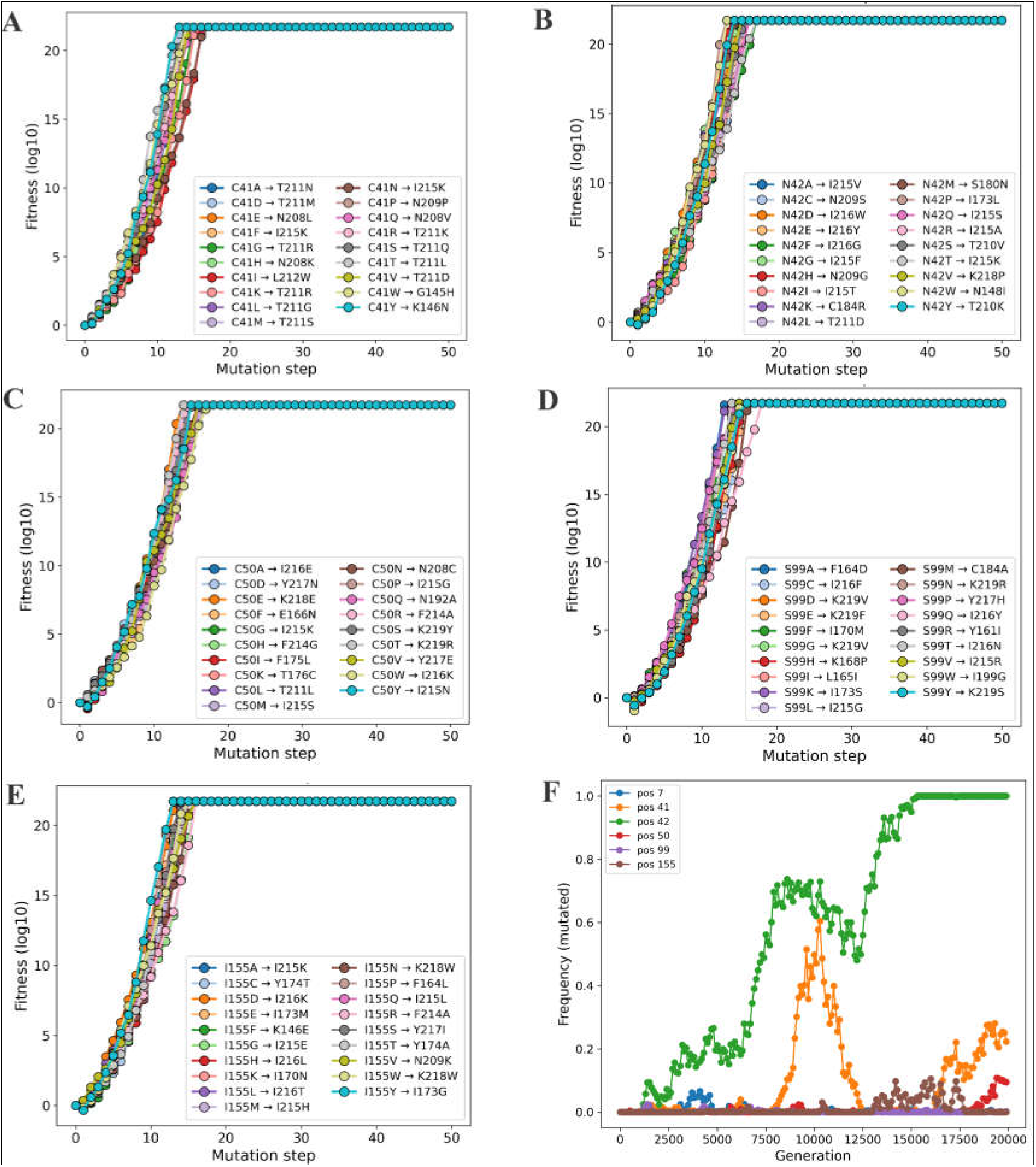
Adaptive-walk trajectories and mutation-frequency dynamics for mutation positions. Panels **A–E** shows stochastic adaptive walks initiated at residues 41, 42, 50, 99 and 155. For each starting position, all 19 amino-acid substitutions rapidly climb toward a common high-fitness plateau but take distinct early paths that converge onto shared compensatory intermediates. (**F**) This shows the population-level mutation frequencies under Wright–Fisher evolution with drug cycling. Position 42 shows the earliest and strongest sweep followed by position 41. Positions 50 and 155 increase only late and weakly. Position 99 remains largely absent and position 7 shows no meaningful selection.

## Discussion

Dihydrofolate reductase has long served as a model for studying adaptive evolution under antifolate pressure (33). Previous studies treated *Pf*DHFR resistance as discrete outcomes with resistance inferred indirectly using surrogates (34, 35). In this work, we combined evolutionary features, network level constraints, molecular dynamics, thermodynamic stability and experimental kinetic data to construct a multidimensional fitness landscape that describes *Pf*DHFR evolution maintaining functionality. Collectively, our findings suggest that *Pf*DHFR is under strong functional constraints. That is, resistance mutations in the binding pocket will only be distinctly tolerated in the presence of compensatory mutations in other regions of the protein (36). The evolutionary and network analyses show that resistance in this protein arises from a collaborative mutational ensemble and not independent substitution.

Co-evolutionary patterns revealed that several mutation sites are statistically coupled forming coordinated clusters rather than acting in isolation. Many of these linked residues fall in or near the binding pocket. This implies that, when one region shifts, other regions somewhere within the protein adjust to maintain overall function. Such coordinated adjustments allow the enzyme to redistribute the structural and dynamic strain, which makes sense of the high-order mutation patterns commonly seen in resistant parasites (35). Structure network and correlation analyses also indicate that resistance mutations rearrange the short-range and long-range communication in *Pf*DHFR. Since these communication pathways link distal sites to the catalytic core, even single substitution can have an indirect effect on the behavior of enzymes by affecting the internal network. Some of these mutations cluster within flexible loop regions that control access to the binding pocket where changes in dynamics influence ligand binding and catalytic activity. Such distal, dynamics-driven effects are consistent with previous studies that have demonstrated long-range allosteric interactions in DHFR (37). Structural correlation maps aided in differentiating between the destabilizing and preservation or restoration of internal connectivity substitutions. The disruptive mutation, N42I, breaks important couplings and discontinue communication channels. This can offer a mechanistic explanation as to why catalytic activity is less as previous studies reported (34, 35). Thus, if a single communication pathway is impaired, alternative pathways emerge to aid in signal transduction, which is an enzymatic property that contributes to resilience. Conversely, other mutations including C41R, C50R, S99N, and I155L do not only destroy the existing contacts, but also form new ones that distant parts of the protein. This type of rearrangement maintains communication over the long range and helps to sustain enzymatic activity. This portrays *Pf*DHFR as a modular system with a stable catalytic core that is supported by adaptable loop regions. Such an architecture could explain why resistance mutations emerge and at the same time preserve sufficient catalytic function. Mutational fitness profiling further emphasizes this modularity, revealing unequal tolerance to substitution across the protein fold. The most conserved residues like 99 and 155 show very little natural variation, yet resistance mutations are reported at these very locations. This apparent contradiction could be explained by epistasis where the effect of a mutation depends on the presence of other mutations (38).

The MD simulations of apo wild-type enzyme and resistant mutants indicate that the resistance mutations do not affect the global fold as was seen by similar values of the RMSD (10, 39). Rather, mutations distribute flexibility to surface-exposed loop regions at active sites and main contact interfaces. This enables the resistant variants to search over a wider number of local conformations and retain the overall structural integrity. Principal components analysis and free-energy analysis indicate that the wild-type enzyme is found mostly in one low-energy basin, and its resistant mutants can freely access multiple low-energy substates separated by relatively small barriers. These flatter energy landscapes provide active flexibility in a direction that does not collapse the conformational ensemble. Prior works have also indicated such flexibility in resistance mutations under selective pressures. Meanwhile, high-order mutations have been associated with reduced conformational flexibility and increased structural rigidity (10). The adaptive simulations via *Pf*PATH showed that antifolate resistance emerges through a restricted set of mutational routes shaped by stability, co-evolution and dynamic couplings. This composite landscape mirrors trends observed in field isolates and laboratory selection experiments (35, 40) in which certain mutations tend to emerge and persist early, followed by the subsequent accumulation of other mutations. Adaptive walks on this landscape reveal that the DHFR protein rarely acquires mutations on their own. Instead, they usually move through a set of residues that act of stabilizing support points before major resistance mutations emerge. These compensatory steps are not confined to the canonical resistance sites but also many other intermediates across the protein fold. These residues form the “latent compensatory ridge” which is a group of structurally permissive positions that buffer destabilizing mutations and at the same time preserve function. Similar observations have been made in DHFR which showed that epistasis and structural constraints restrict viable adaptive routes yielding reproducible mutational trajectories across populations (41, 42). All in all, *Pf*DHFR adaptation relies not only on active-site chemistry but also on the broader architectural framework that distributes mechanical strain and tunes conformational ensembles.

The simulations at population level also indicate a hierarchical order of replacements which agrees with patterns observed in natural *P. falciparum* populations. The allele-frequency patterns indicated a clear hierarchy of substitutions. Position 42 (51) rose first and most rapidly, followed by a delayed rise in position 41 (50). Positions 50 (59), 99 (108) and 155 (164) were almost flat indicating their great catalytic cost in the composite landscape. Through this ordering, the overall trend of natural *P. falciparum* populations is observed with the DHFR mutations generally accumulating in the sequence of N51I -C59R -S108N -I164L, N51I (IRNL) (10, 40). Although S108N is often described as the first resistance mutation in natural populations (10), this haplotype in our experiment stayed at low frequency across generations. Under the fitness landscape and drug-cycling regime used, stronger early gains were consistently realized at positions 41 and 42. Hence, it is obvious that the simulations favored substitutions that balance resistance with catalytic viability and penalize destabilizing changes unless supported by the compensatory ridge. This could explain why some resistance sites are highly accessible and others remain conditional or rare. It could also explain why natural parasites repeatedly follow similar evolutionary routes and why only a small subset of all possible DHFR mutational combinations is observed in field isolates.

Our study elucidates how DHFR in *Plasmodium falciparum* adapts under antifolate pressure by evolving along a narrow and constrained evolutionary path. This trajectory allows the enzyme to balance the fitness cost of resistance while retaining sufficient catalytic activity for survival. Nonetheless, this study has limitations. Our analyses focused on the apo form of *Pf*DHFR and did not explicitly consider ligand- or drug-bound states. Despite these limitations, the strong agreement between evolutionary features, structural dynamics and experimentally established resistance patterns supports a mechanistic view of *Pf*DHFR adaptation to antifolate pressure. Crucially, the presence of a narrow compensatory ridge suggests that resistance evolution in this protein is restricted to a limited set of structurally permissive pathways. Experimentally targeting this latent ridge may offer a rational strategy to both validate these predictions and guide the design of antifolates that constrain the parasite’s ability to access viable escape routes.

## Materials and Methods

### Sequence preprocessing

For this study, we used the crystal structure of the wild-type *P. falciparum* DHFR-TS protein (PDB ID: 3QGT) as the reference for our analyses. The protein structure consists of two domains: the DHFR domain (first 231 residues) and the TS domain (last 288 residues), separated by an 89-residue interjunction region (Fig. S2). Given the focus on DHFR, we truncated the DHFR domain, which stretches from amino acid position 10 to 228 in the full-length protein as annotated in UNIPROT (43). This truncation was performed using the PYMOL Molecular Graphics System Version 3.0.3 (44). For consistency, the truncated sequence was re-indexed from 1 to 219, a scheme used throughout this study.

### Homology Modeling and Structure Validation

Upon examination, the DHFR protein contained ten missing residues in its sequence from position 86 to 95. To fill in these gaps, homology modeling was performed using Modeller (Version 10.5) (45, 46). This software generated several protein models and ranked them using the Discrete Optimized Protein Energy (DOPE) scoring function. The model with the lowest DOPE score was selected as the most accurate representation of the full DHFR structure (Fig. S2). To ensure the accuracy of the modeled structure, we calculated the Root Mean Square Deviation (RMSD) between the modeled residues and the crystal structure (Fig. S3). An RMSD value of 0.306Å indicated high structural similarity. After validation, we extracted the amino acid sequence of length 219 (as shown below) from the modeled DHFR structure for further analyses. The modeled structure also served as the seed structure for MD simulations.

>A7UD81_PLAFA

DIYAICACCKVESKNEGKKNEVFNNYTFRGLGNKGVLPWKCNSLDMKYFCAVTTYVNESKYEKLKYKRCKYLNKETVDNVNDMPNSKKLQNVVVMGRTSWESIPKKFKPLSNRINVILSRTLKKEDFDEDVYIINKVEDLIVLLGKLNYYKCFIIGGSVVYQEFLEKKLIKKIYFTRINSTYECDVFFPEINENEYQIISVSDVYTSNNTTLDFIIYK K

### Sequence Space Analysis

To explore the mutational landscape of DHFR, we performed a sequence space analysis using the multiple sequence alignment (MSA) of homologous DHFR proteins. First, we retrieved 5000 non-redundant protein sequences homologous to DHFR using the NCBI BlastP tool (47). The sequences were cleaned to ensure high quality, non-redundancy and biological relevance. We then aligned the cleaned sequences using the ClustalW algorithm (48) and the resulting MSA served as the basis for the subsequent analyses.

### Occupancy Calculation

Occupancy calculations were performed to assess how frequently specific amino acids occupy each residue position in homologous DHFR sequences (49). Positions with high occupancy imply conserved residues which are likely essential for the stability of protein. We identified residues where mutations may have significant impacts on the protein’s structure and function by comparing occupancy data with mutation sites.

### Shannon Entropy Calculation

Shannon entropy (S (*i*)) (49, 50) was calculated to quantify the variability of amino acids at each sequence position. Positions with low entropy values represent conserved residues, critical for structural and functional integrity, while high entropy values indicate positions prone to mutations. Shannon entropy was calculated following this formulation,

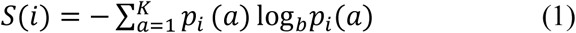

Where ***i*** represents the sequence position, *a* indexes the amino-acid alphabet of size *K* (with *K*=20 for standard protein sequences) and *p_i_*(*a*) is the probability of observing amino acid *a* at position *i* in the multiple sequence alignment. Lower values of *S*(*i*) correspond to highly conserved residues, while higher values indicate increased sequence variability.

### Mutual Information Calculation

To investigate co-evolving residues, we calculated mutual information (MI) (49, 51, 52) for each pair of positions in the MSA. Mutual information quantifies the degree of correlation between two sequence positions, revealing residues that might be functionally or structurally interdependent. These co-varying residues could indicate regions where mutational compensation occurs, maintaining overall protein function despite mutations in specific positions.

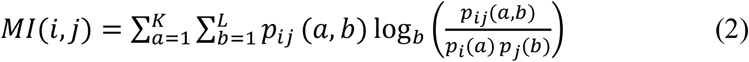

In this formulation, *a* and *b* index the amino-acid alphabets at positions *i* and *j*, of sizes *K* and *L*, respectively (with *K*=*L*=20 for protein sequences). Here, *p_ij_*(*a*,*b*) denotes the joint probability of observing amino acids *a* and *b* at positions *i* and *j*, while *p_i_*(*a*) and *p_j_*(*b*) represent the corresponding marginal probabilities at each position. The MI *(i,j)* values range from 0 (uncorrelated residues) to MI*_max_* (the most interdependent residue pairs). In this study, a cutoff MI value of 0.5 was chosen to focus on highly co-varying positions. The MI values were visualized as a heatmap and a co-evolutionary matrix, generated using python programming.

### Intrinsic Disorder Prediction

Intrinsic disorder prediction analysis was conducted to explore the presence of disordered regions within the DHFR protein. Disordered regions, known for their inherent flexibility, can tolerate higher levels of mutation and often play significant roles in protein-protein interactions. For a comprehensive evaluation, we employed six widely used disorder predictors: PONDR® FIT (21), PONDR® VSL2 (22), PONDR® VL3 (22), PONDR® VLXT (23), IUPred Short (24), and IUPred Long (24). These predictors use distinct predictive approaches to identify disordered regions within the protein. PONDR® VSL2 is particularly optimized for proteins containing both structured and disordered regions (53). We combined the results of all the six predictors to obtain an average disorder profile (MDP), which yielded a unanimous disorder score to each residue. The scores above 0.5 were considered as disordered and the ones between 0.15 and 0.5 as flexible but considered relatively ordered. This disorder profile was further analyzed to look at possible correlations between disordered regions, known mutation sites, and active site residues to help understand the role of protein flexibility in resistance mutations and the maintenance of protein function.

### Sequence-based Network Analysis

Following the MI calculations, we performed a sequence-based network analysis to identify co-evolving residues. Nodes in the network represent individual residues, while edges denote mutual information-based connections between them. MI thresholds were set to define the strength of these interactions, with different ranges indicating very strong (MI ≥ 0.8), strong (0.6 < MI < 0.8), moderate (0.4 < MI < 0.6) and weak (0.0 < MI < 0.4) interactions. This helped us visualize a network of interactions for selected residues of interest, particularly mutation sites and active site residues. The resulting networks provided insight into how mutations at specific positions might influence stability and drug resistance through interdependence with nearby residues. To investigate densely connected sub-networks within the protein, we employed Maximal Clique Analysis (MCA) (25, 26) and Community Detection (27) methods. In MCA, we aimed to identify groups of residues that form maximal cliques, where every residue within a clique is directly connected to every other residue. This method focuses on identifying highly interconnected sets of residues that are structurally and functionally interdependent. Each residue in the protein is represented as a node and the edges between these nodes represent significant interactions, which are based on proximity in the protein structure or known co-evolutionary relationships. Community network analysis was also performed to divide the protein’s residue interaction network into distinct subgroups or communities. Unlike MCA, which focuses on fully connected sub-networks, community detection identifies groups of residues that are more likely to interact within their community than with residues outside it.

### Structure Network and Correlation Analysis

Dynamic correlation and network analysis of *Pf*DHFR were done with the Bio3D package (version 2.4-1.9000) of R (30, 54). Bio3D offers a platform to comparative structure analysis, normal mode analysis and analysis of MD-based correlation network. Minimization of the protein structures of the wild-type and mutant PfDHFR variants was initially performed followed by alignment of the backbone atoms to achieve the same frames of reference. In our case, we used the dccm() function of Bio3D to calculate residue-residue cross-correlation matrices (C*ij*) to an extent that quantifies concerted atomic motions of individual structures. These matrices were then offered as inputs to correlation network analysis through the cna() function which represents each residue as a node and then pairs with weighted edges between the nodes based on how they move with each other. We used a cut (of |C*ij*| ≥ 0.3) to remove weak correlations and concentrate on couplings that have a functional meaning. The resulting networks were partitioned into communities with the Girvan-Newman edge betweenness algorithm (55) which represents clusters of highly correlated residues that are associated with dynamically consistent modules. The visualizations of networks were created with the plot.cna() and pymol() functions and examined in PyMOL 3D representations of the community of residues and communication pathways. In-house Python/R scripts were used to do other visualization and customized correlation mapping to create dynamic cross-correlation maps and network overlays.

### Molecular Dynamic Simulations and Trajectory Analysis

A 1 microsecond MD simulation of apo *Pf*DHFR was performed to examine the structural dynamics using GROMACS 2025.1 (56) for both wild type and mutant structures. The system was prepared with the Amber99SB-ILDN force field (57) that provides superior side-chain torsion parameters to proteins. The *Pf*DHFR structure was put in a cubic periodic box (side length of around 8.5 nm) and solvated with 25, 476 explicit SPC/E water molecules (58). Neutralizing counter-ions (8 Cl⁻) were added to mimic a 0.15 M ionic strength environment and long-range electrostatic interactions were treated with the particle-mesh Ewald (PME) method (59, 60). The system was first energy minimized until convergence (< 1000 kJ/mol/nm maximum force) was achieved before production. The equilibration was then carried out in two stages, that is, 100ps NVT at 300 K and 100ps NPT at 1 bar. A 2fs integration timestep was used, with all covalent bonds involving hydrogen constrained via the LINCS algorithm (61) to stabilize the system. Energies and pressures were stored as well as trajectory coordinates every 10ps and every 1ps respectively, to be analyzed later. During the production run, we undertook the standard structural descriptors to describe the dynamic activity of the protein. The backbone root-mean-square deviation was computed to determine the general conformational drift and equilibration. Per-residue root-mean-square fluctuations were determined to determine the flexible regions of the protein. The DSSP algorithm was also used to track secondary structure content and thus monitor any alterations in helices, 2-sheet or loops (62). The other descriptors that were considered were the radius of gyration, solvent-accessible surface area, intra-molecular hydrogen bonds and principal component analysis to obtain the major motions and the major conformational states. The first two major components projected the free energy landscape to show the relative stabilities of the various conformational substates. These analyses presented in totality a global view of the conformational dynamics of PfDHFR throughout the simulation.

### Simulating Evolutionary Trajectories in Heterogeneous Landscapes

We developed Plasmodium Fitness Pathway Analysis of Trajectories in Heterogeneous Landscapes (*Pf*PATH) to investigate how *Plasmodium falciparum* DHFR adapts under antifolate pressure. *Pf*PATH is a population-genetic and adaptive-walk simulator for *Pf*DHFR designed to computationally analyze how the protein adapts under drug pressure, structural constraints and mutational noise in high-dimensional heterogeneous fitness landscapes. This computational model integrated different layers of sequence information, structure features and experimentally derived kinetic and drug-response constraints into a unified fitness landscape and propagates parasite genotypes through a Wright-Fisher evolutionary process (32). The conceptual model is a generalization of the biophysical trajectory models already used to study DHFR evolution in other systems (34), except that it captures aspects of evolution, structural constraints and experimentally measured drug-response behavior. All genotypes were represented as full DHFR amino-acid sequences with mutations defined relative to the wild-type (WT). For a genotype *g*, the set of substitutions indexes

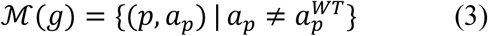

the positions and amino acids mutated from WT. Each mutation was assigned a set of single-site scores reflecting functional impact, stability changes and structure-based energetic effects. Since these sources differed in scale, each score was converted to a dimensionless quantity by z-normalization and hyperbolic-tangent transformation,

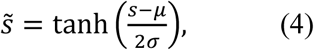

which scaled down the data to the range of −1 to 1 so that no individual dataset dominated the terrain. Stability-related quantities were inverted in sign such that stabilizing mutations made a positive contribution. The composite score of genotype *g* was calculated by summing the contributions of all mutated sites. The single-site effects represented tendencies in functionality as inferred from mutational scanning and the anticipated alterations in thermodynamic stability. Pairwise terms were then added to the model to approximate epistasis (41). These consisted of co-evolutionary couplings and structural–dynamics couplings. A small positive contribution was assigned when both members of a strongly interacting pair were mutated, whereas a small penalty was imposed when only one member of the pair was mutated. Multi-mutant ΔG predictions were also incorporated into the model. This allowed the landscape to capture non-additive mutation–mutation interactions. These relations rendered the terrain rugged and directional. To restrict exploration to biologically feasible mutational loads, *Pf*PATH applied a mild penalty to genotypes containing more than six substitutions. This cutoff was used to reflect the rarity of *Pf*DHFR haplotypes with more than six mutations in nature. Genotypes containing more than six mutations were assigned a linearly proportional mutation-load penalty per additional substitution. An explicit stability filter based on ΔΔG predictions was also applied. For each genotype, the most destabilizing predicted ΔΔG was identified as,

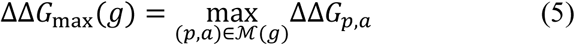

A smooth penalty was applied whenever this value exceeded a soft threshold, and any genotype whose maximum ΔΔG exceeded a hard cutoff was excluded. This stability gate prevented evolutionary paths from passing through highly unstable or catalytically incompetent intermediates. The final composite score was

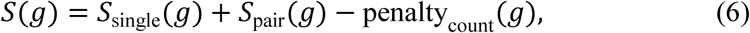

and intrinsic fitness is mapped via a Boltzmann-like transformation,

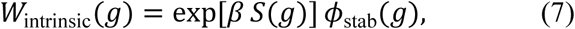

with selection strength *β* = 2. This formulation ensured that moderate differences in composite score translated into meaningful but not extreme differences in reproductive success. Drug pressure was applied by adjusting intrinsic fitness based on standard DHFR resistance haplotypes. The mutations N51I, C59R, S108N, and I164L were used to classify genotypes as wild-type-like, double-, triple-, or quadruple-resistant. These resistant haplotypes were assigned a multiplicative fitness benefit in the presence of drug, proportional to experimentally measured increases in inhibitor dissociation constants. In the absence of drug, these haplotypes incurred a fitness cost corresponding to reduced catalytic efficiency, as previously reported for these mutants (40). During drug-on periods,

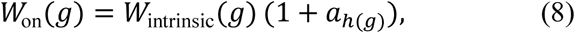

and in drug-off periods,

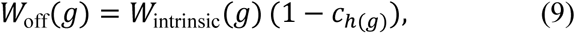

where *a_h_*_(_*_g_*_)_ and *c_h_*_(_*_g_*_)_ depend on the haplotype class. Simulations assumed either constant drug pressure or cyclical exposure, allowing exploration of how intermittent treatment reshaped evolutionary trajectories. Evolution in *Pf*PATH was modeled using a Wright–Fisher framework, which describes a population with discrete, non-overlapping generations in which each generation is formed by sampling individuals from the previous generation according to relative fitness (63). The population size *N* was fixed at 10,000 individuals per generation, and offspring were sampled using multinomial sampling with replacement. Mutations occurred independently at each amino-acid site with probability *μ* per generation. Although natural *P. falciparum* mutation rates are low, an elevated effective mutation rate (*μ* = 10⁻⁵ per site) was used to capture both de novo mutation and standing variation, enabling multi-step adaptive paths to be observed on computationally feasible timescales. After mutation, each unique genotype was assigned a fitness value using the composite fitness scorer, and the next generation of *N* individuals was sampled proportionally to these fitness values. Allele and haplotype frequencies at selected positions of interest were tracked across generations to capture time-resolved evolutionary dynamics in the presence and absence of drug.

## Acknowledgements

DM acknowledges One Health Trust, Bangalore, India. DM and SC acknowledge the Director, Birla Institute of Technology and Science-Pilani, Hyderabad, India.

## Author Contributions

Conceptualization: SC and DS. Methodology: DM. Formal analysis: DM. Figure Preparation: DM and VNU. Writing—original draft: DM. Writing—review & editing: SC, DS, VNU, PU, and DP.

## Data and materials availability

All data needed to evaluate the conclusions in the paper are present in the paper and/or the Supplementary Materials.

