## Supplementary Material for "Adaptive Landscapes of *Plasmodium Falciparum* Dihydrofolate Reductase Reveal Pathways to Antifolate Resistance"

**Figures**

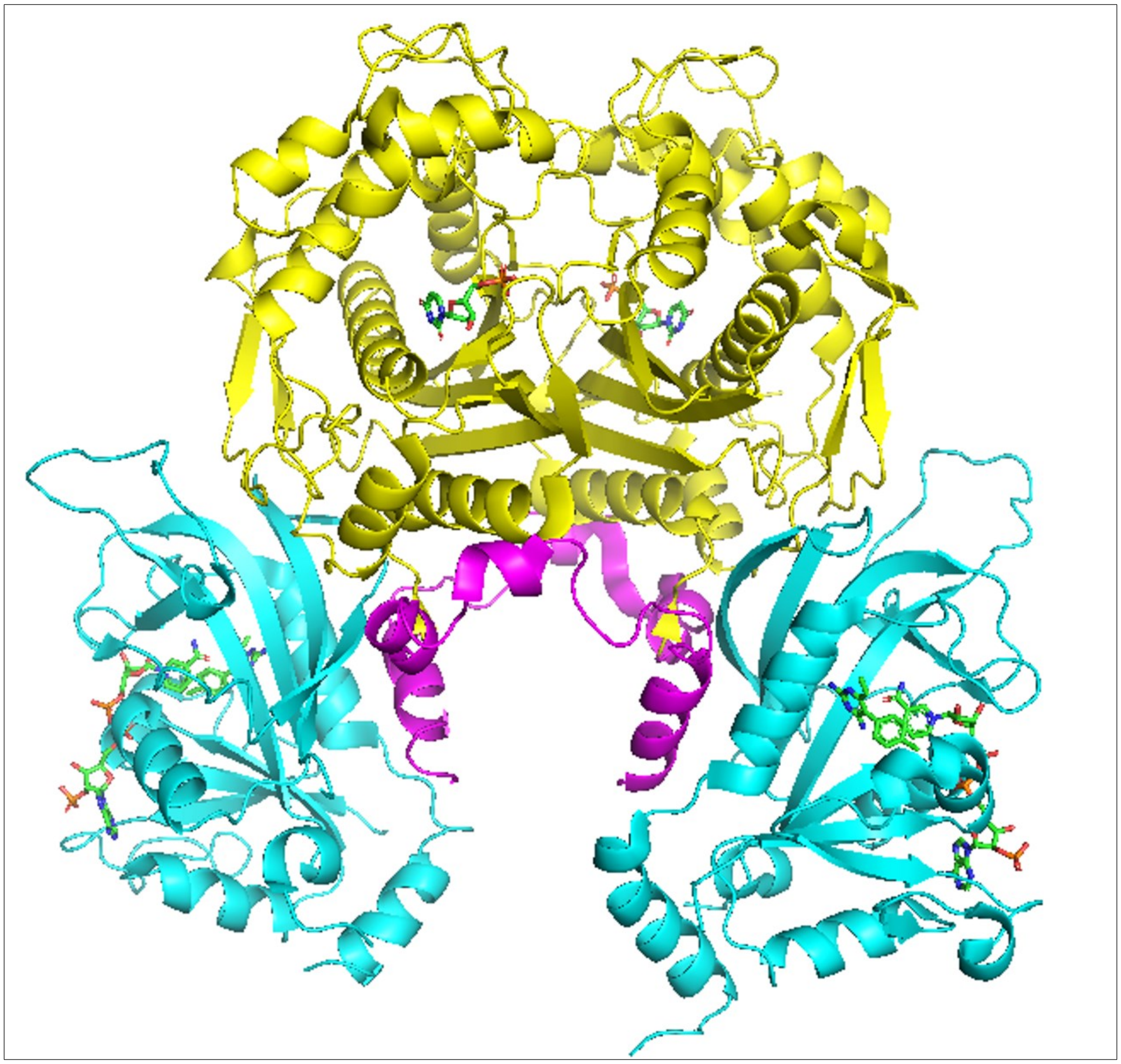

**Fig. S1. This illustrates the 608 long pfDHFR-TS (PDB ID: 3QGT) dimeric assembly**. In cyan are the DHFR domains with DHFR-TS junction in magenta and yellow region as TS domains. The DHFR domains are bound with the drug pyrimethamine and NADPH cofactor.

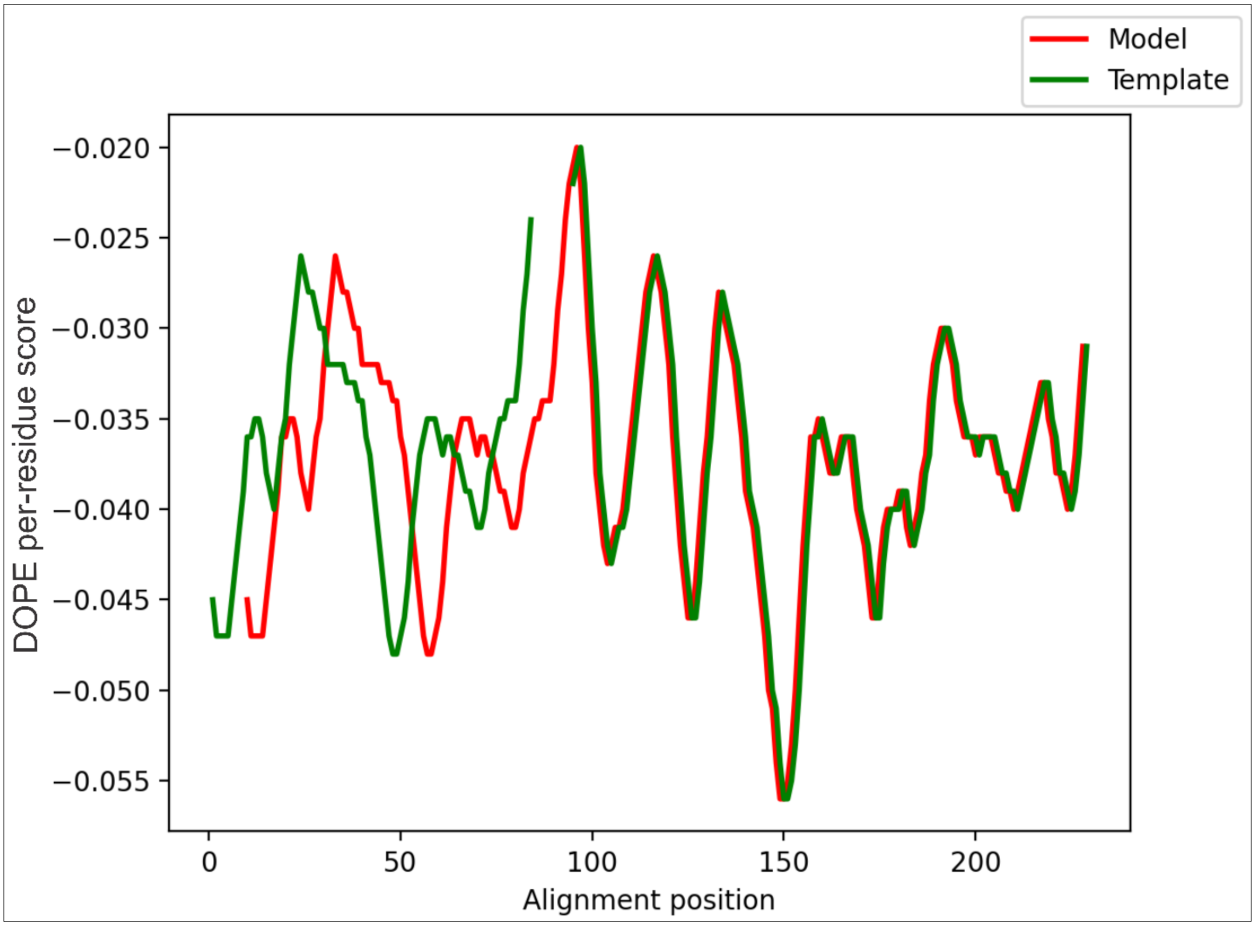


**Fig. S2. The DOPE scores for the model and template structures.** This shows the validation of homology modeling that was done with the use of Modeller. A comparison is made between the DOPE scores in the alignment positions. The correlation of the model (red) and the template (green) is depicted in the plot. However, the misalignment observed between positions 0 to 96 is expected due to the missing residues gap in the template structure.

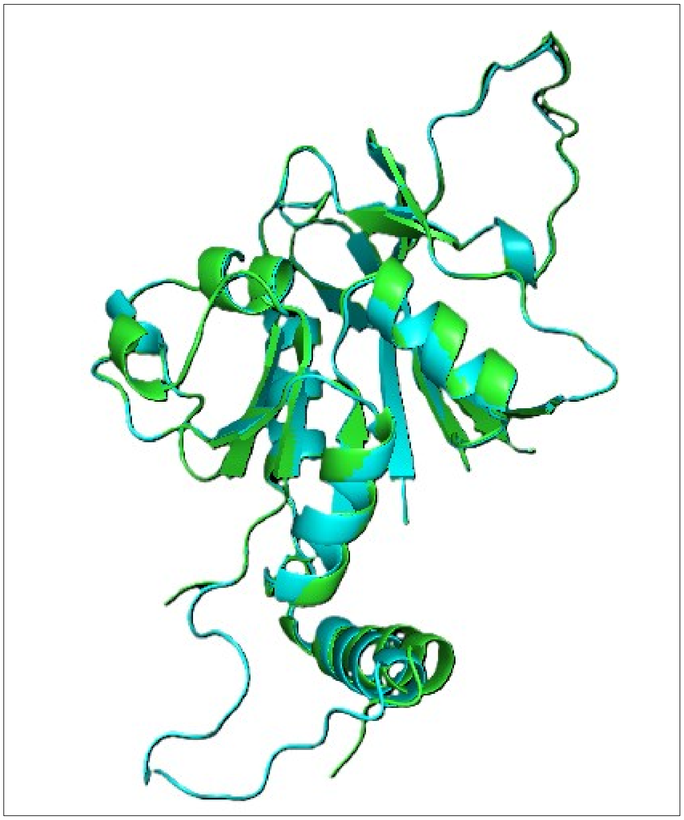

**Fig. S3. The structural alignment of the modeled DHFR**. This depicts the DHFR domain (in cyan) with the template structure (in green) using PyMOL. The RMSD was calculated to assess the structural similarity between the model and the template. The RMSD value of 0.306 Å indicates a good alignment, validating the accuracy of the homology model. This alignment helps confirm that the modeled structure retains the key features and overall topology of the template.

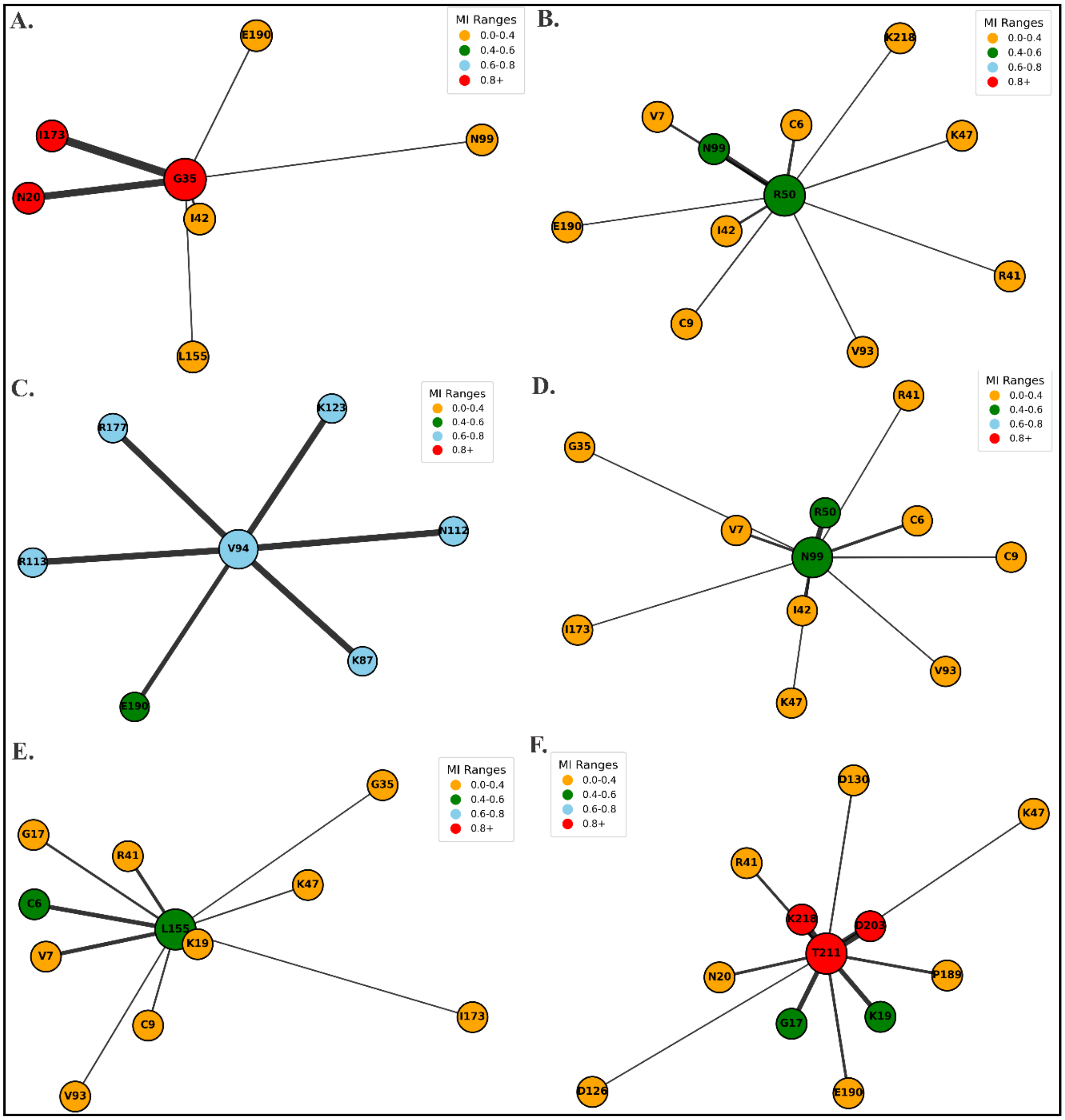


**Fig. S4. Mutational interaction networks reveal both local and distal sub-networks that overlap with active-site communication paths.** (Related to Fig. 2). (**A**) G35 forms a small short-range cluster comprising of I42, N20, I173 and N99, which shows tight local communication. (**B-D**) These networks (R50 and N99) reveal some common connections between mutation sites (41, 42, 50, 99) and active residues nearby (C6, V7, R41), which indicates that these locations may form hubs of interaction between catalytic and mutation-prone regions. (**E**) Position 155, a mutation–active site, shows intermediate-range contacts with C6, K19 and R41. (**F**) Long-range coupling around T211 links spatially distant residues D203 and K19 to earlier loop positions G17 and R41. These networks show how mutation-prone residues are wired into both local and distal sub-networks that overlap with active-site communication paths.

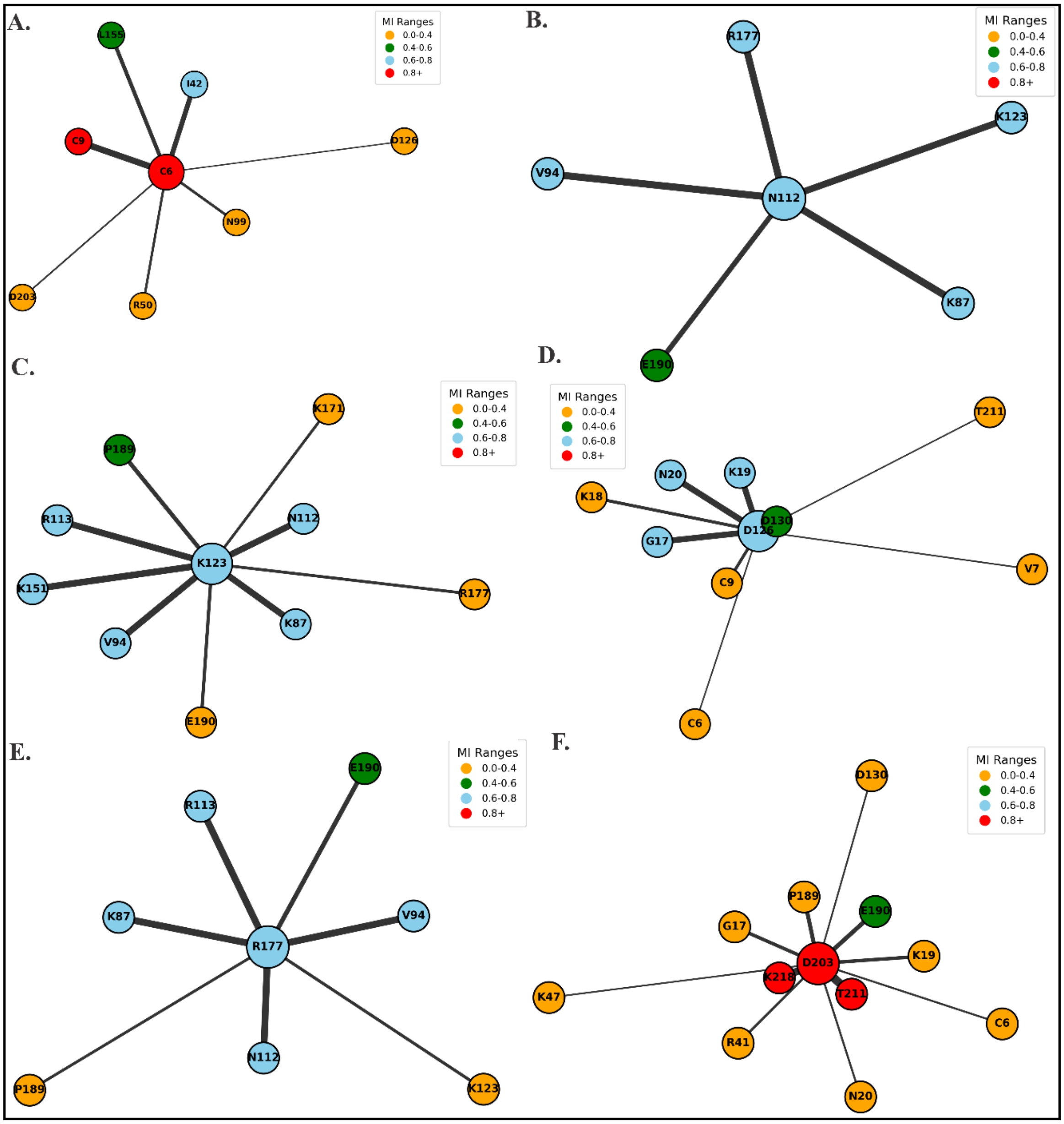


**Fig. S5. Interaction networks highlighting tightly connected sub-networks with strong MI couplings.** (Related to Fig. 2). These networks place emphasis on a group of residues that repeatedly appear in well-coordinated compact clusters throughout the protein. A lot of these residues interact with each other strongly at a short and long range and tend to connect loops, strands, and adjacent structural blocks. These clusters have lots of nodes that contain known sites of mutations or residues that continuously co-occur with the mutation hotspots in other analysis. They are highly connected indicating that they might serve as small stabilizing sub-modules that cushion or re-distribute local perturbations, which may offer possible compensatory benefits in the event of mutations in local regions.

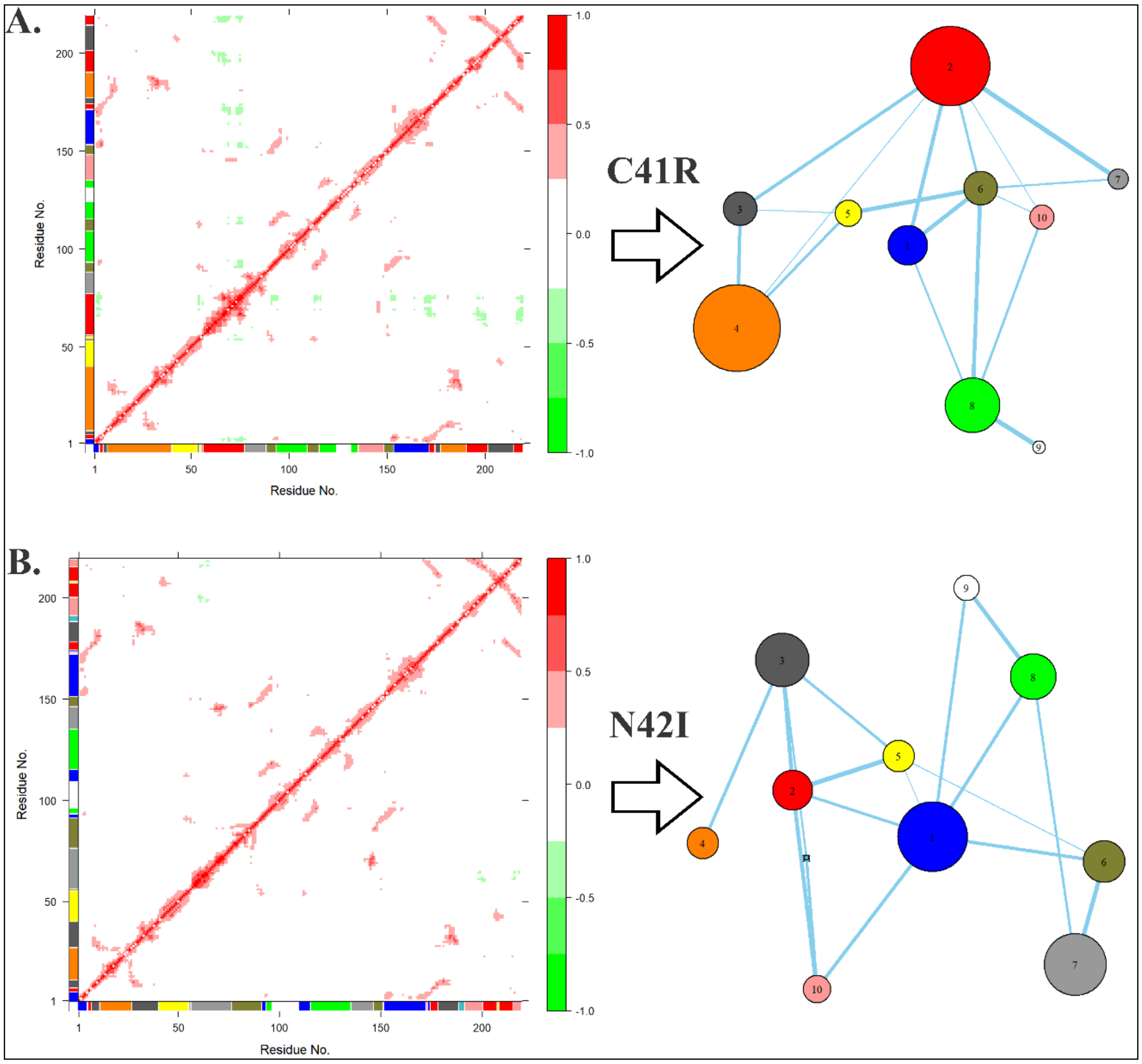


**Fig. S6. Local flexibility and correlated motions in in mutants C41R and N42I.** (Related to Fig. 4). (**A**) C41R DCCM experiences a decrease in positive-correlation patches which are seen in the WT. We however observe more anticorrelated movements occur in the fold. The community network insert reveals a reorganization into a larger set of communities. C41R lies in a block (Cluster 5) with another catalytic residue C50. (**B**) The N42I DCCM displays strengthened correlations in the N-terminal segment and mild weakening in distal regions. The insert on the right indicates that N42I remains in the same structural block containing residues 41-55.

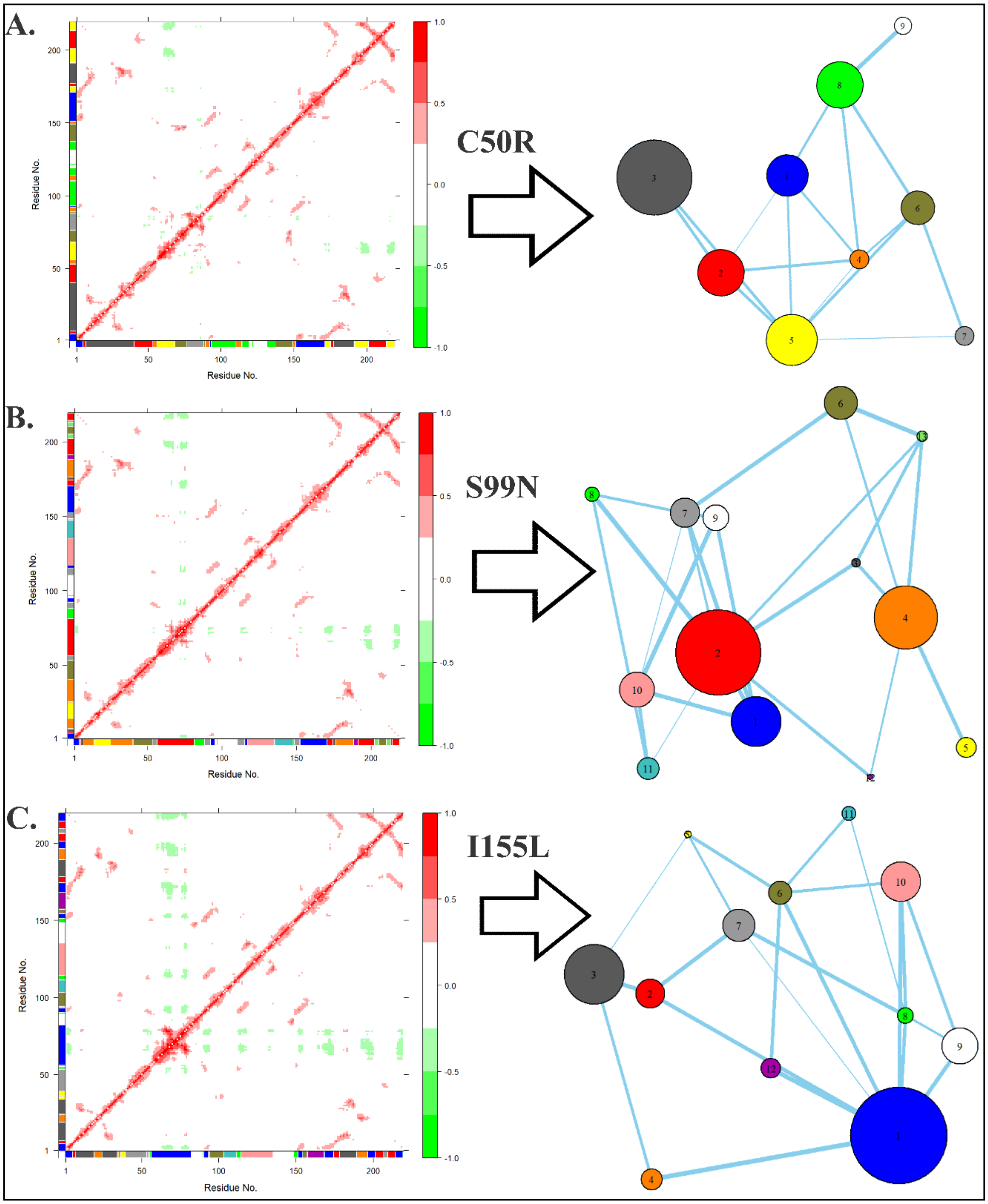


**Fig. S7. Local flexibility and correlated motions in in mutants C50R, S99N and I155L.** (Related to Fig. 4). (**A**) C50R shows broader anticorrelated regions compared to WT with fewer strong long-range contacts. The community network is smaller and C50 reorganization is rearranged into other modules. (**B**) S99N loses several correlations toward the C-terminal region and gains scattered anticorrelated patches. The community map shows residues around 99–103 clustering together in a small module (**C**) I155L displays stronger correlations around the central 68–79 region and new anticorrelated zones along the fold. The community network redistributes several residues, with I155 joining the cluster that typically contains 99, 102 and 103.

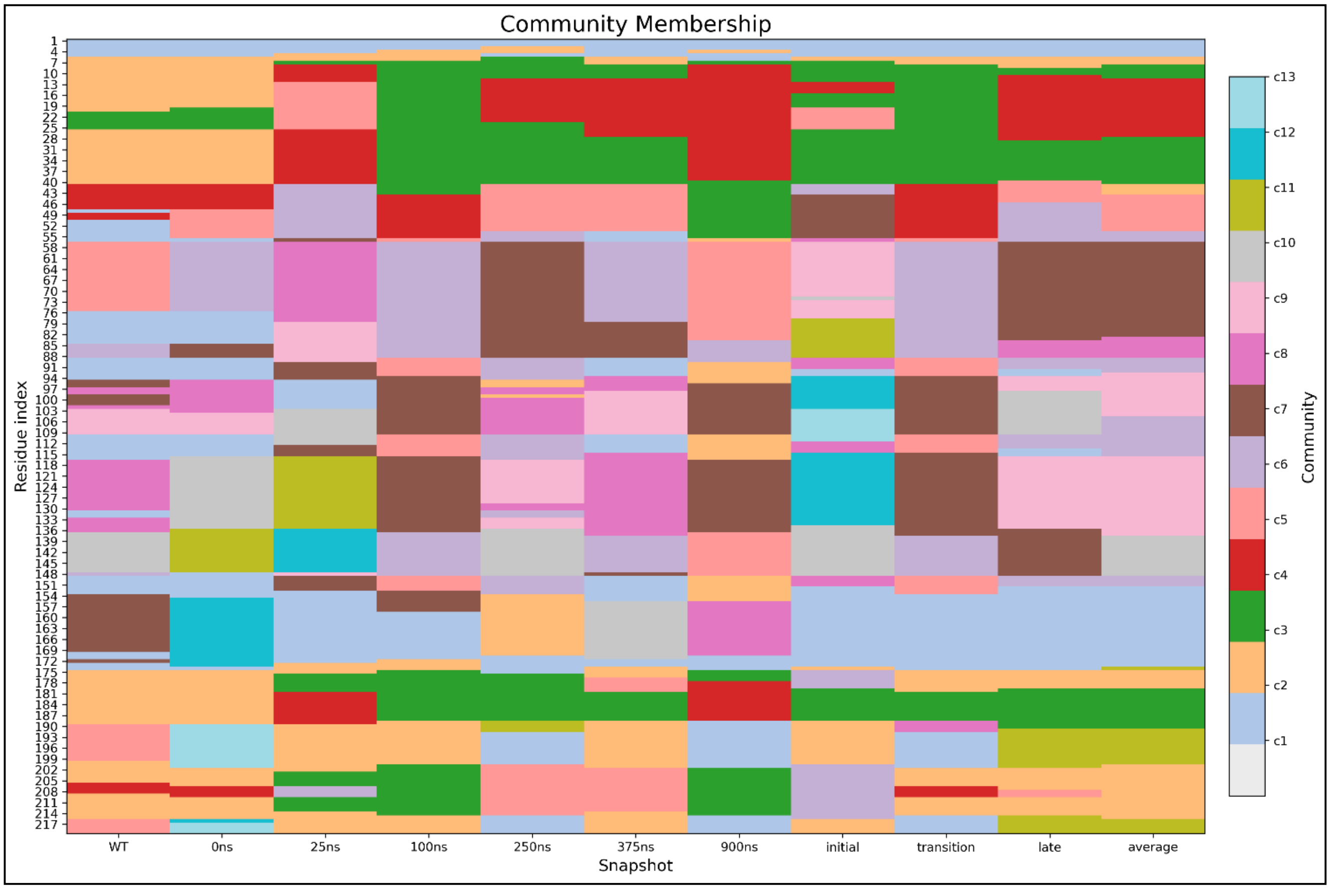


**Figure S8. Community membership changes across the WT PfDHFR trajectory.** (Related to Fig. 4 and Fig. 5). This follows the dynamics of each residue changing community membership between the significant snapshots of the WT MD simulation. The seed WT structure is positioned in the first column and the snapshots at 0ns, 25ns, 100ns, 250ns, 375ns and 900ns follow in that order. The final four columns show averaged snapshots for the early state ("initial"), the transition window ("transition"), the late metastate ("late"), and the overall average across 1µns ("average"). The residues that persist in the same community in the snapshots manifest as a continuous block of the same hue, while residues that shift communities show patch-like transitions. The N-terminal area is quite stable and has the same community along the trajectory. However, many internal and loop regions change membership along the trajectory. The implication of these shifts is that the local reorganization of correlated motions that occur during the transition does not interfere with the global communication structure of the WT enzyme.

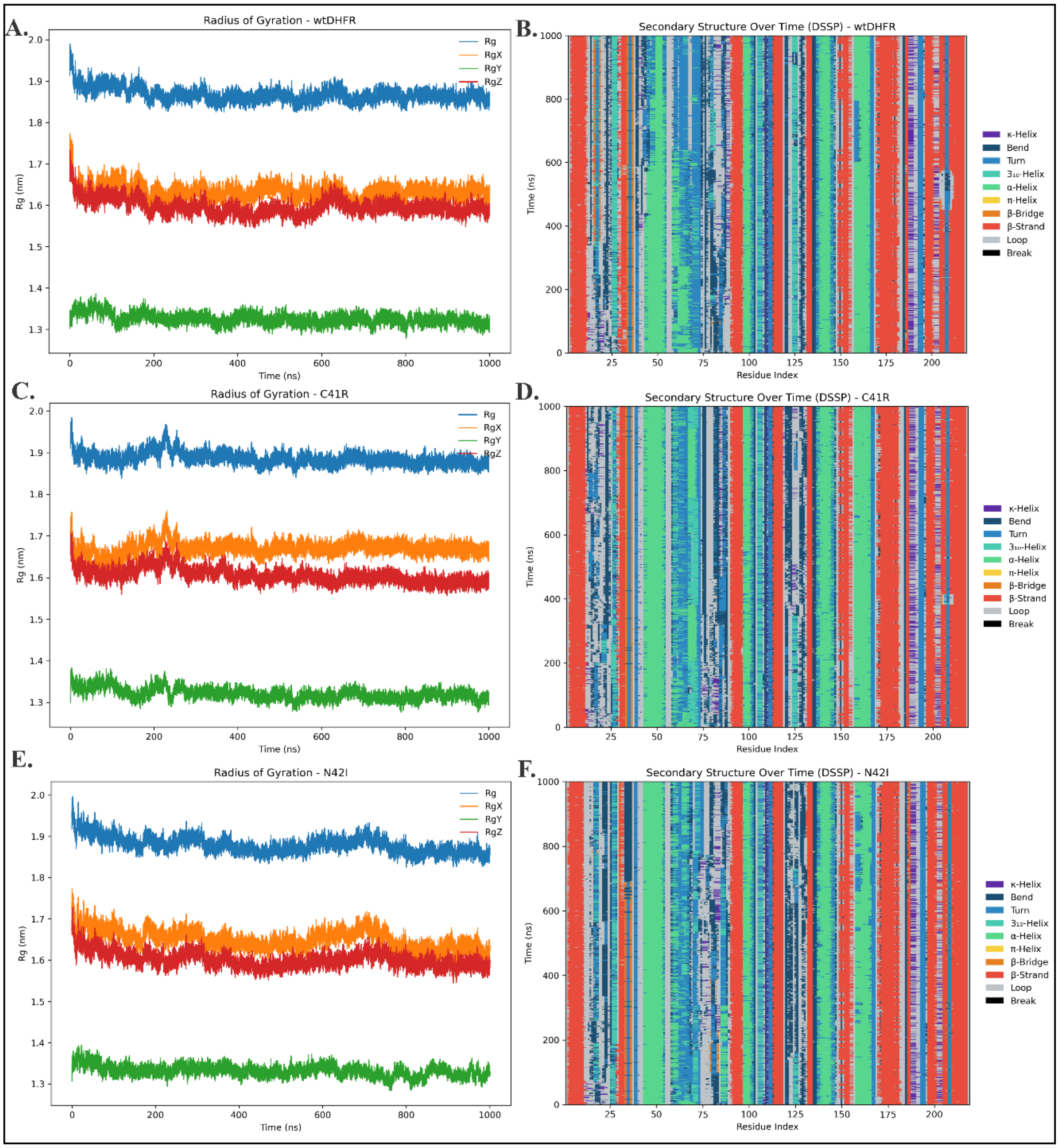


**Figure S9. Radius of gyration and secondary-structure stability of WT and mutants**. (Related to Fig. 5). (**A, C, E**) The overall Rg (blue trace) does not change significantly between the WT and the two mutants over the 1 µs simulation. This indicates that the global fold stays compact and does not undergo large-scale expansion. The RgX, RgY and RgZ are variable around constant baselines, indicating rather natural directional breathing patterns than structural drift. Mutants show slightly wider fluctuations than the WT, but no major deviations, which is consistent with a global architecture being preserved. (**B, D, F**) The DSSP profiles indicate that the main secondary-structure components, namely, α-helices and β-strands, are not lost during the course of the simulation in both mutants and the WT. The greatest variation is in loop and turn regions that inherently vary and are more dynamic in the mutants. There are some instances of transition between bends, turns and loops but they do not show development or loss of core structure.

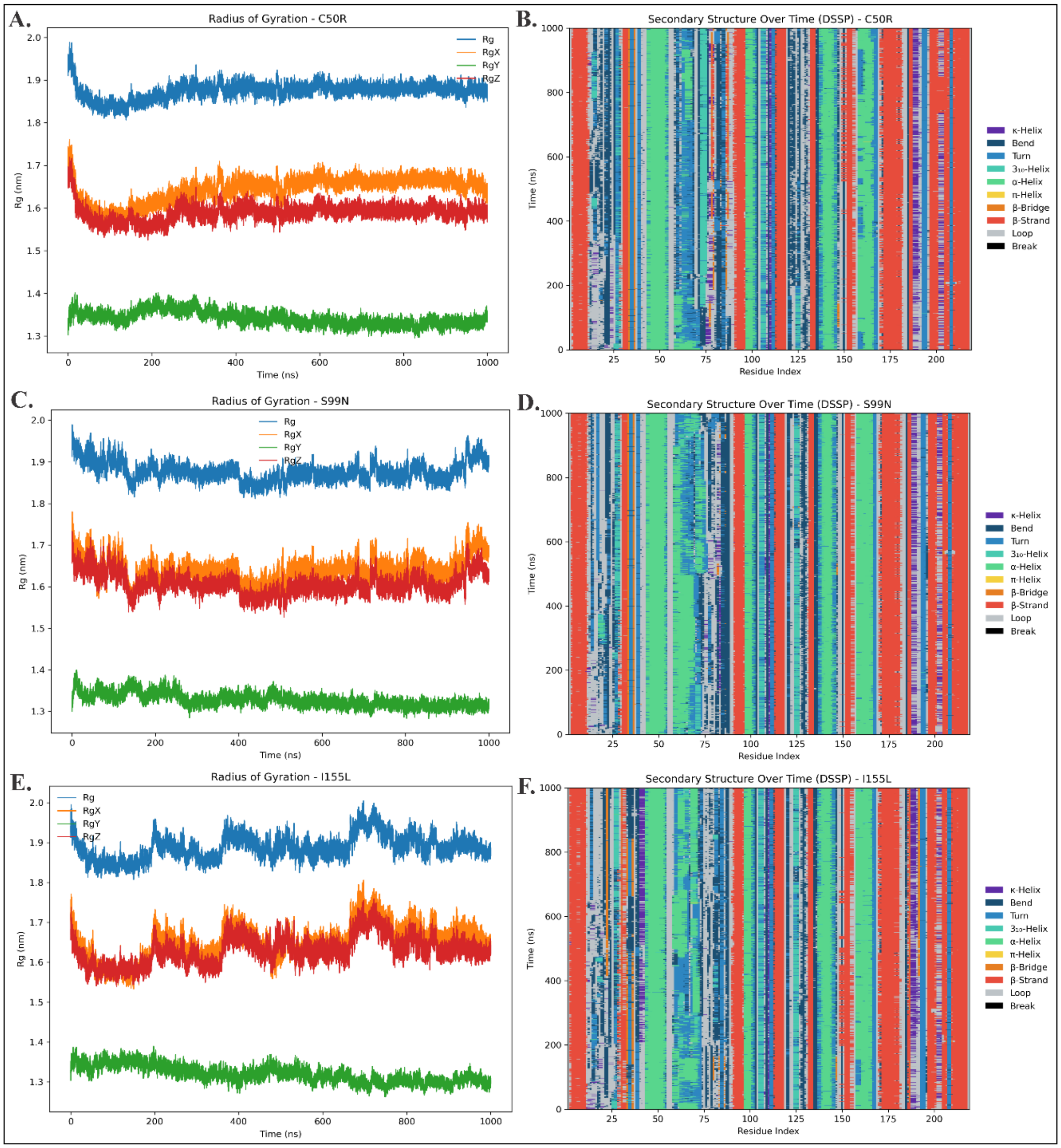


**Figure S10. Radius of gyration and secondary-structure stability for additional *Pf*DHFR mutants**. (Related to Fig. 5). (**A, C, E**) For all three mutants, the overall Rg remains steady throughout the trajectory. The directional components (RgX, RgY, RgZ) fluctuate within stable ranges reflecting normal breathing motions rather than structural drift. The mutants exhibit a bit higher amplitude variations than the WT, which is in line with a higher local flexibility yet global compactness has been maintained. (**B, D, F**) Across the mutants, the α-helices and β-strands mostly remain intact for the entire simulation. The majority of structural changes take place in loop and turn regions that naturally change and are more mobile in the mutants. There is no significant loss of secondary structure. This implies that the main architecture is held constant when there is flexibility change locally.

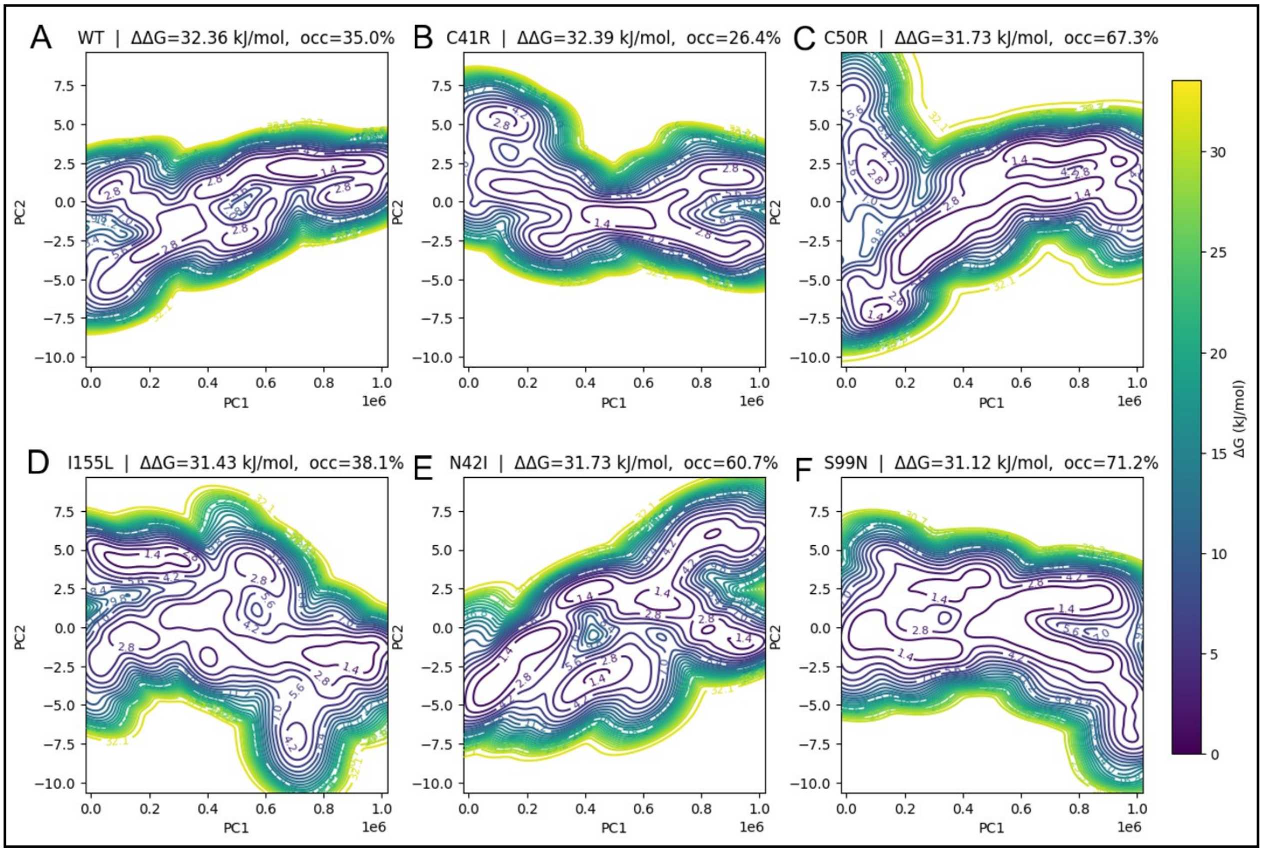


**Fig. S11. Free-energy landscapes (apo PfDHFR) reveal a rugged WT funnel and progressively flattened mutant ensembles with comparable global energy spans.** (Related to Fig. 6). (**A)** WT exhibits a steep, rugged funnel with multiple ridges separating deep minima. This is an indication of constrained conformational breathing around a dominant folded basin. (**B**) C41R exhibits a broader saddle-like landscape with several shallow dimples. (**C**) C50R shows a wide, near-degenerate basin. This may allow facile transitions among substates. (**D**) I155L forms a broad primary basin with low-barrier satellites. (**E**) N42I reveals a gently corrugated trough. (**F**) S99N demonstrates the flattest profile. It is characterized by a series of similar minima which are associated with loop flexibility.

**Tables**

**Table S1:** Variable *Pf*DHFR residue positions identified from the *P. falciparum*–specific Shannon entropy analysis. (Related to Fig. 1A).

| Position | Residue | Shannon Entropy | Residue Name |
|---|---|---|---|
| 6 | C | 0.135794 | Cysteine |
| 7 | A | 0.132691 | Alanine |
| 9 | C | 0.124251 | Cysteine |
| 17 | G | 0.116907 | Glycine |
| 18 | K | 0.404114 | Lysine |
| 20 | N | 0.116907 | Asparagine |
| 35 | G | 0.110453 | Glycine |
| 41 | C | 0.110453 | Cysteine |
| 42 | N | 0.660060 | Asparagine |
| 47 | K | 0.110453 | Lysine |
| 50 | C | 0.480439 | Cysteine |
| 87 | K | 0.110453 | Lysine |
| 94 | V | 0.110453 | Valine |
| 99 | S | 0.480439 | Serine |
| 112 | N | 0.110453 | Asparagine |
| 113 | R | 0.110453 | Arginine |
| 123 | K | 0.276381 | Lysine |
| 126 | D | 0.188113 | Aspartic acid |
| 151 | K | 0.110453 | Lysine |
| 155 | I | 0.542349 | Isoleucine |
| 171 | K | 0.276381 | Lysine |
| 173 | I | 0.110453 | Isoleucine |
| 177 | R | 0.224451 | Arginine |
| 189 | P | 0.139061 | Proline |
| 190 | E | 0.158411 | Glutamic acid |
| 203 | D | 0.271189 | Aspartic acid |
| 211 | T | 0.304636 | Threonine |
| 218 | K | 0.410116 | Lysine |

**Table S2:** A summary of entropy values for conserved and functionally relevant residues derived from the broader DHFR homolog alignment across multiple organisms (Related to Fig. 1A)

| Position | Residue | Shannon Entropy | Residue Name |
|---|---|---|---|
| 30 | G | 0 | Glycine |
| 32 | G | 0 | Glycine |
| 38 | P | 0 | Proline |
| 39 | W | 0 | Tryptophan |
| 91 | N | 0 | Asparagine |
| 104 | P | 0 | Proline |
| 37 | L | 0.036434 | Leucine |
| 176 | T | 0.064581 | Threonine |
| 45 | D | 0.089578 | Aspartic acid |
| 46 | M | 0.317875 | Methionine |
| 155 | I | 0.371818 | Isoleucine |
| 48 | Y | 0.374846 | Tyrosine |
| 49 | F | 0.400231 | Phenylalanine |
| 5 | I | 0.444473 | Isoleucine |
| 7 | A | 0.492124 | Alanine |
| 6 | C | 0.662822 | Cysteine |
| 47 | K | 0.729842 | Lysine |
| 42 | N | 0.901884 | Asparagine |
| 41 | C | 0.963004 | Cysteine |
| 99 | S | 1.239797 | Serine |
| 50 | C | 1.646328 | Cysteine |

**Table S3:** Key co-varying residues associated with each of the selected mutation sites showing the mutual-information correlations and Cα–Cα structural distances (Related to Fig. 1B)

| Mutation Site | Co-varying residues | MI Correlation | Distance from Mutation site |
|---|---|---|---|
| A7 | **41** | **0.803** | **11.274** |
| | **42** | **0.488** | **10.041** |
| | **50** | 0.146 | 12.626 |
| | **99** | 0.185 | 12.992 |
| | 126 | 0.016 | 28.524 |
| | **155** | **0.391** | **9.513** |
| | 218 | 0.040 | 16.280 |
| R41 | **7** | **0.803** | **11.274** |
| | **50** | 0.0014 | 13.563 |
| | **99** | 0.008 | 14.260 |
| | **155** | **0.267** | **15.800** |
| | 211 | 0.185 | 13.418 |
| | 203 | 0.141 | 20.003 |
| | 218 | 0.361 | 27.049 |
| N42 | **6** | **0.632** | **12.746** |
| | **7** | **0.488** | **10.041** |
| | 9 | 0.207 | 11.354 |
| | 20 | 0.029 | 27.493 |
| | 35 | 0.134 | 16.053 |
| | 47 | 0.183 | 8.488 |
| | **50** | 0.159 | 12.105 |
| | 93 | 0.050 | 19.577 |
| | **99** | 0.174 | 16.160 |
| | 151 | 0.006 | 24.204 |
| | 173 | 0.134 | 19.197 |

**Table S4.** Community membership in the wild-type structure (Related to Fig. 4D)

| id | size | members |
|---|---|---|
| 1 | 35 | c(1:5, 56, 76:84, 89:94, 110:115, 148:154, 174) |
| 2 | 55 | c(6:19, 26:40, 175:189, 202:206, 210:215) |
| 3 | 6 | 20:25 |
| 4 | 10 | c(41:47, 207:209) |
| 5 | 8 | 48:55 |
| 6 | 19 | 57:75 |
| 7 | 4 | 85:88 |
| 8 | 9 | 95:103 |
| 9 | 6 | 104:109 |
| 10 | 20 | 116:135 |
| 11 | 12 | 136:147 |
| 12 | 20 | c(155:173, 216) |
| 13 | 15 | c(190:201, 217:219) |

**Table S5.** Community membership of the mutant structure C41R (Related to Fig. 4)

| id | size | members |
|---|---|---|
| 1 | 24 | c(1, 54:56, 77:84, 89:92, 110:113, 149:152) |
| 2 | 24 | c(2:5, 93, 153:171) |
| 3 | 57 | c(6:40, 176:189, 204:206, 210:214) |
| 4 | 8 | c(41:45, 207:209) |
| 5 | 8 | 46:53 |
| 6 | 20 | 57:76 |
| 7 | 4 | 85:88 |
| 8 | 32 | c(94:103, 114:135) |
| 9 | 6 | 104:109 |
| 10 | 13 | 136:148 |
| 11 | 23 | c(172:175, 190:203, 215:219) |

**Table S6.** Community membership of the mutant structure N42I (Related to Fig. 4)

| id | size | members |
|---|---|---|
| 1 | 32 | c(1, 54:56, 77:84, 89:92, 108:116, 131:132, 148:152) |
| 2 | 25 | c(2:5, 93:94, 153:171) |
| 3 | 61 | c(6:19, 26:42, 175:190, 202:215) |
| 4 | 6 | 20:25 |
| 5 | 11 | 43:53 |
| 6 | 20 | 57:76 |
| 7 | 4 | 85:88 |
| 8 | 13 | 95:107 |
| 9 | 18 | c(117:130, 133:136) |
| 10 | 11 | 137:147 |
| 11 | 18 | c(172:174, 191:201, 216:219) |

**Table S7.** Community membership of the mutant structure C50R (Related to Fig. 4)

| id | size | members |
|---|---|---|
| 1 | 39 | c(1:5, 54:56, 77:84, 89:94, 110:116, 131:132, 149:155, 172) |
| 2 | 44 | c(6:19, 26:40, 176:190) |
| 3 | 6 | 20:25 |
| 4 | 18 | c(41:44, 46:47, 203:214) |
| 5 | 7 | c(45, 48:53) |
| 6 | 21 | c(57:76, 218) |
| 7 | 4 | 85:88 |
| 8 | 9 | 95:103 |
| 9 | 6 | 104:109 |
| 10 | 17 | c(117:130, 133:135) |
| 11 | 13 | 136:148 |
| 12 | 19 | c(156:171, 173:175) |
| 13 | 16 | c(191:202, 215:217, 219) |

**Table S8.** Community membership of the mutant structure S99N (Related to Fig. 4)

| id | size | members |
|---|---|---|
| 1 | 46 | c(1, 56, 77:84, 89:94, 110:135, 149:152) |
| 2 | 27 | c(2:5, 153:174, 216) |
| 3 | 50 | c(6:12, 17:18, 26:40, 175:189, 202:207, 210:214) |
| 4 | 11 | c(13:16, 19:25) |
| 5 | 8 | c(41:45, 48, 208:209) |
| 6 | 9 | c(46:47, 49:55) |
| 7 | 20 | 57:76 |
| 8 | 4 | 85:88 |
| 9 | 8 | 95:102 |
| 10 | 7 | 103:109 |
| 11 | 13 | 136:148 |
| 12 | 16 | c(190:201, 215, 217:219) |

**Table S9.** Community membership of the mutant structure I155L (Related to Fig. 4)

| id | size | members |
|---|---|---|
| 1 | 32 | c(1, 54:56, 77:84, 89:92, 108:116, 131:132, 148:152) |
| 2 | 25 | c(2:5, 93:94, 153:171) |
| 3 | 61 | c(6:19, 26:42, 175:190, 202:215) |
| 4 | 6 | 20:25 |
| 5 | 11 | 43:53 |
| 6 | 20 | 57:76 |
| 7 | 4 | 85:88 |
| 8 | 13 | 95:107 |
| 9 | 18 | c(117:130, 133:136) |
| 10 | 11 | 137:147 |
| 11 | 18 | c(172:174, 191:201, 216:219) |

**Table S10.** Summary of Maximal Clique Analysis of the *Pf*DHFR (Related to Fig. 3A)

**Table S10.1. Largest Maximal Cliques Detected**

| Rank | Clique ID | Clique Size | Member Residues |
|---|---|---|---|
| 1 | - | 11 | 78, 79, 80, 81, 91, 92, 93, 113, 114, 115, 152 |
| 2 | - | 10 | 11, 30, 31, 32, 33, 38, 183, 184, 185, 186 |
| 3 | 3 | 10 | 30, 31, 32, 33, 37, 38, 183, 184, 185, 186 |
| 4 | - | 10 | 78, 79, 80, 81, 92, 93, 113, 114, 115, 116 |
| 5 | - | 10 | 78, 79, 80, 91, 92, 93, 94, 114, 115, 152 |

**Table S10.2. Cliques Containing Catalytic (Active-Site) Residues**

| Clique ID | Size | Active Residues Present |
|---|---|---|
| 3 | 10 | 37 |
| 6 | 9 | 7 |
| 7 | 9 | 7 |
| 8 | 9 | 7, 176 |
| 9 | 9 | 7, 176 |
| 24 | 9 | 37 |
| 25 | 9 | 37 |
| 34–37 | 8 | 7 |
| 43 | 8 | 176 |
| 52–54 | 8 | 37 |
| 64–66 | 8 | 161 |
| 75 | 7 | 155 |
| 81 | 7 | 176 |
| 83 | 7 | 37 |
| 95–97 | 7 | 161 |
| 101–102 | 7 | 176 |
| 114 | 6 | 5, 155 |
| 115 | 6 | 5 |
| 122 | 6 | 37 |
| 124 | 6 | 45, 46 |
| 125 | 6 | 45, 46, 48 |
| 126 | 6 | 46, 48, 49 |
| 127–128 | 6 | 48, 49 |
| 141, 143–144 | 6 | 155 |
| 145–147 | 6 | 155, 161 |
| 148 | 6 | 99, 102 |
| 167–168 | 5 | 5 |
| 169–171 | 5 | 7, 176 |
| 176 | 5 | 45, 46, 48, 49 |
| 189 | 5 | 155 |
| 193, 195–196 | 5 | 99 |
| 198 | 5 | 99, 102, 103 |
| 215, 218 | 5 | 176 |

| **Clique ID** | **Size** | **Active Residues Present** |
|---|---|---|
| 223 | 4 | 5, 6 |
| 224 | 4 | 6, 176 |
| 225 | 4 | 6, 7, 176 |
| 234 | 4 | 103 |
| 255 | 3 | 102, 103 |

**Table S10.3. Cliques Containing Mutation-Prone Residues**

| Clique ID | Size | Mutation Residues Present |
|---|---|---|
| 6–9 | 9 | 7 |
| 34–37 | 8 | 7 |
| 75 | 7 | 155 |
| 114 | 6 | 155 |
| 123–124 | 6 | 42 |
| 126–128 | 6 | 50 |
| 141, 143–144, 146–147 | 6 | 155 |
| 148 | 6 | 99 |
| 169–171 | 5 | 7 |
| 175 | 5 | 42 |
| 177 | 5 | 50 |
| 189 | 5 | 155 |
| 193, 195–196, 198 | 5 | 99 |
| 225 | 4 | 7 |
| 229 | 4 | 41, 42 |
| 252 | 3 | 41 |

**Table S10.4. Critical Cliques Containing Both Catalytic and Mutation Residues**

| Clique ID | Size | Active Residues | Mutation Residues |
|---|---|---|---|
| 6 | 9 | 7 | 7 |
| 7 | 9 | 7 | 7 |
| 8 | 9 | 7, 176 | 7 |
| 9 | 9 | 7, 176 | 7 |
| 34–37 | 8 | 7 | 7 |
| 75 | 7 | 155 | 155 |
| 114 | 6 | 5, 155 | 155 |
| 124 | 6 | 45, 46 | 42 |
| 126 | 6 | 46, 48, 49 | 50 |
| 127–128 | 6 | 48, 49 | 50 |
| 141 | 6 | 155 | 155 |
| 143–144 | 6 | 155 | 155 |
| 146–147 | 6 | 155, 161 | 155 |
| 148 | 6 | 99, 102 | 99 |
| 169–171 | 5 | 7, 176 | 7 |
| 189 | 5 | 155 | 155 |
| 193, 195–196 | 5 | 99 | 99 |
| 198 | 5 | 99, 102, 103 | 99 |
| 225 | 4 | 6, 7, 176 | 7 |

**Table S11.** Community network analysis of PfDHFR showing nine communities, each representing a group of residues that move cohesively within the protein's dynamic architecture (Related to Fig. 3B)

| ID | Size | Active-Site Residues | Mutation-Prone Residues | Communities with Both Active + Mutation Sites |
|---|---|---|---|---|
| 0 | 15 | 99, 102, 103 | 99 | Yes |
| 1 | 33 | - | - | No |
| 2 | 31 | 37 | - | No |
| 3 | 30 | 6, 7, 176 | 7 | Yes |
| 4 | 13 | 45, 46, 48, 49 | 41, 42, 50 | Yes |
| 5 | 15 | 161 | - | No |
| 6 | 19 | - | - | No |
| 7 | 40 | 5, 155 | 155 | Yes |
| 8 | 23 | - | - | No |

**Table S12.** Tabulation of intrinsic disorder prediction scores for PfDHFR using different PONDR algorithms. (Related to Fig. 1B).

| Index pos. | Res | VLXT | XL1_XT | CAN_XT | VL3 | VSL2 |
|---|---|---|---|---|---|---|
| 1 | D | 0.002311339 | 0 | 0 | 0.3253 | 0.376499 |
| 2 | I | 0.004830018 | 0 | 0 | 0.3138 | 0.317174 |
| 3 | Y | 0.007496831 | 0 | 0 | 0.304 | 0.277501 |
| 4 | A | 0.011870756 | 0 | 0 | 0.2916 | 0.198309 |
| 5 | I | 0.053839753 | 0 | 0 | 0.2819 | 0.157827 |
| 6 | C | 0.07629971 | 0 | 0 | 0.2741 | 0.112303 |
| 7 | A | 0.092293986 | 0 | 0 | 0.2672 | 0.091732 |
| 8 | C | 0.113397262 | 0 | 0 | 0.2642 | 0.111294 |
| 9 | C | 0.136352297 | 0 | 0 | 0.2643 | 0.137397 |
| 10 | K | 0.170839547 | 0 | 0 | 0.2649 | 0.217574 |
| 11 | V | 0.167663253 | 0 | 0 | 0.2695 | 0.328241 |
| 12 | E | 0.164955723 | 0 | 0 | 0.2773 | 0.449806 |
| 13 | S | 0.152758966 | 0 | 0 | 0.283 | 0.542904 |
| 14 | K | 0.126421207 | 0 | 0 | 0.2854 | 0.639281 |
| 15 | N | 0.104748456 | 1.42E-08 | 4.50E-07 | 0.2875 | 0.703444 |
| 16 | E | 0.090461075 | 0.003209573 | 5.90E-07 | 0.2837 | 0.741358 |
| 17 | G | 0.072713506 | 0.048847502 | 5.90E-07 | 0.2652 | 0.707829 |
| 18 | K | 0.053742392 | 0.096539446 | 5.91E-07 | 0.2461 | 0.738874 |
| 19 | K | 0.013754183 | 0.173977986 | 5.90E-07 | 0.2327 | 0.72585 |
| 20 | N | 0.013981557 | 0.255831918 | 1.42E-07 | 0.2153 | 0.649948 |
| 21 | E | 0.014582564 | 0.337822406 | 1.42E-07 | 0.197 | 0.568705 |
| 22 | V | 0.014997249 | 0.421101804 | 1.42E-07 | 0.1823 | 0.472004 |
| 23 | F | 0.014985784 | 0.504345452 | 1.42E-07 | 0.17 | 0.370903 |
| 24 | N | 0.014009358 | 0.586248997 | 1.42E-07 | 0.1611 | 0.296411 |

| | | | | | | |
|---|---|---|---|---|---|---|
| 25 | N | 0.011772385 | 0.663298651 | 1.30E-09 | 0.1534 | 0.247733 |
| 26 | Y | 0.007590286 | 0.683480951 | 1.30E-09 | 0.1456 | 0.21379 |
| 27 | T | 0.002290484 | 0.635789041 | 0 | 0.1403 | 0.200891 |
| 28 | F | 0.001529275 | 0.558350506 | 0 | 0.1348 | 0.198271 |
| 29 | R | 0.00126179 | 0.476496576 | 0 | 0.1296 | 0.222069 |
| 30 | G | 0.000700294 | 0.39450609 | 0 | 0.1228 | 0.224872 |
| 31 | L | 0.000243376 | 0.311226692 | 0 | 0.1149 | 0.232145 |
| 32 | G | 0.000227405 | 0.227983045 | 0 | 0.1104 | 0.238202 |
| 33 | N | 0.004231722 | 0.146079554 | 7.00E-10 | 0.1076 | 0.229432 |
| 34 | K | 0.004400864 | 0.065820343 | 7.00E-10 | 0.1072 | 0.218618 |
| 35 | G | 0.004570006 | 1.37E-07 | 7.00E-10 | 0.1103 | 0.21041 |
| 36 | V | 0.004571452 | 1.02E-07 | 7.00E-10 | 0.1099 | 0.197217 |
| 37 | L | 0.004570298 | 9.48E-08 | 7.00E-10 | 0.1088 | 0.19089 |
| 38 | P | 0.004567547 | 9.03E-08 | 7.00E-10 | 0.108 | 0.191866 |
| 39 | W | 0.004498963 | 8.75E-08 | 7.00E-10 | 0.1067 | 0.178772 |
| 40 | K | 0.004475185 | 8.39E-08 | 7.00E-10 | 0.1022 | 0.170287 |
| 41 | C | 0.004445753 | 8.21E-08 | 7.00E-10 | 0.0983 | 0.159802 |
| 42 | N | 0.000374014 | 2.64E-08 | 0 | 0.0912 | 0.150112 |
| 43 | S | 0.000198504 | 2.52E-08 | 0 | 0.0824 | 0.134531 |
| 44 | L | 2.35E-05 | 1.00E-10 | 0 | 0.0756 | 0.124851 |
| 45 | D | 1.95E-05 | 1.00E-10 | 0 | 0.0759 | 0.124577 |
| 46 | M | 2.33E-05 | 0 | 0 | 0.0752 | 0.119363 |
| 47 | K | 2.66E-05 | 0 | 0 | 0.0809 | 0.106381 |
| 48 | Y | 0.000489364 | 0.000001795 | 0 | 0.0852 | 0.100507 |
| 49 | F | 0.000779633 | 9.00E-06 | 0 | 0.0917 | 0.093535 |
| 50 | C | 0.000929177 | 1.83E-05 | 0 | 0.1004 | 0.077777 |
| 51 | A | 0.001067879 | 2.08E-05 | 0 | 0.112 | 0.084777 |

| | | | | | | |
|---|---|---|---|---|---|---|
| 52 | V | 0.001099082 | 0.012108016 | 0 | 0.1257 | 0.093633 |
| 53 | T | 0.001189573 | 0.012146962 | 0 | 0.1402 | 0.115728 |
| 54 | T | 0.00152921 | 0.012146965 | 0 | 0.1554 | 0.158452 |
| 55 | Y | 0.0016358 | 0.012146966 | 0 | 0.1732 | 0.201571 |
| 56 | V | 0.006194728 | 0.012146968 | 0 | 0.1902 | 0.273603 |
| 57 | N | 0.013219439 | 0.012145176 | 0 | 0.2077 | 0.343067 |
| 58 | E | 0.018734762 | 0.012137976 | 0 | 0.2251 | 0.409207 |
| 59 | S | 0.025292335 | 0.012128707 | 0 | 0.2423 | 0.45767 |
| 60 | K | 0.075833355 | 0.012126185 | 0 | 0.2591 | 0.499205 |
| 61 | Y | 0.152029224 | 3.90E-05 | 0 | 0.2775 | 0.507524 |
| 62 | E | 0.219098073 | 0.00000002 | 0 | 0.2977 | 0.524472 |
| 63 | K | 0.285917944 | 1.72E-08 | 0 | 0.3167 | 0.521409 |
| 64 | L | 0.388993681 | 1.62E-08 | 0 | 0.336 | 0.518172 |
| 65 | K | 0.463944024 | 1.42E-08 | 0 | 0.3496 | 0.509862 |
| 66 | Y | 0.527578658 | 0.000000011 | 0 | 0.3599 | 0.477625 |
| 67 | K | 0.544302106 | 8.10E-09 | 0 | 0.3755 | 0.453278 |
| 68 | R | 0.609401872 | 5.80E-09 | 0 | 0.388 | 0.433971 |
| 69 | C | 0.59767269 | 5.30E-09 | 0 | 0.3984 | 0.414575 |
| 70 | K | 0.527917974 | 5.60E-09 | 0 | 0.4114 | 0.413668 |
| 71 | Y | 0.470131139 | 7.43E-08 | 0 | 0.4253 | 0.413105 |
| 72 | L | 0.405840373 | 5.24E-07 | 0 | 0.4369 | 0.441473 |
| 73 | N | 0.332800385 | 0.029000088 | 0 | 0.4486 | 0.439798 |
| 74 | K | 0.283433256 | 0.068633384 | 0.00000023 | 0.4591 | 0.461784 |
| 75 | E | 0.212546159 | 0.143019726 | 0.000134886 | 0.4695 | 0.499455 |
| 76 | T | 0.19066666 | 0.227464983 | 0.000177341 | 0.4743 | 0.525052 |
| 77 | V | 0.120538323 | 0.312211942 | 0.000177347 | 0.4826 | 0.533563 |
| 78 | D | 0.091326045 | 0.396948795 | 0.091723221 | 0.4921 | 0.563357 |

| | | | | | | |
|---|---|---|---|---|---|---|
| 79 | N | 0.096539046 | 0.505839693 | 0.185879101 | 0.503 | 0.602229 |
| 80 | V | 0.09613359 | 0.616950731 | 0.279846822 | 0.5117 | 0.618527 |
| 81 | N | 0.12667912 | 0.728059093 | 0.374886436 | 0.5173 | 0.628339 |
| 82 | D | 0.118032173 | 0.810167093 | 0.470087362 | 0.5213 | 0.63169 |
| 83 | M | 0.115820958 | 0.88163945 | 0.565310327 | 0.5225 | 0.63022 |
| 84 | P | 0.153709092 | 0.918341467 | 0.659783187 | 0.5256 | 0.637321 |
| 85 | N | 0.22817568 | 0.932512882 | 0.770851843 | 0.5294 | 0.635188 |
| 86 | S | 0.284171646 | 0.888134785 | 0.859350398 | 0.5317 | 0.633344 |
| 87 | K | 0.289810976 | 0.886025357 | 0.871062101 | 0.5322 | 0.621575 |
| 88 | K | 0.299898326 | 0.848457889 | 0.853810257 | 0.5376 | 0.603239 |
| 89 | L | 0.330222997 | 0.819157638 | 0.760682207 | 0.5464 | 0.576721 |
| 90 | Q | 0.35783102 | 0.790141983 | 0.665647056 | 0.5541 | 0.547366 |
| 91 | N | 0.418460797 | 0.71323382 | 0.587824151 | 0.5596 | 0.485496 |
| 92 | V | 0.498295114 | 0.659936792 | 0.513436721 | 0.5558 | 0.445422 |
| 93 | V | 0.502720618 | 0.625607508 | 0.49526294 | 0.5463 | 0.420576 |
| 94 | V | 0.470153113 | 0.612047902 | 0.435213679 | 0.5343 | 0.394567 |
| 95 | M | 0.419737816 | 0.657406631 | 0.401216706 | 0.524 | 0.409951 |
| 96 | G | 0.411619516 | 0.664868059 | 0.298167061 | 0.5159 | 0.403418 |
| 97 | R | 0.398413979 | 0.690832947 | 0.235352295 | 0.5055 | 0.425344 |
| 98 | T | 0.367688952 | 0.647143931 | 0.31878571 | 0.4918 | 0.418551 |
| 99 | S | 0.371937445 | 0.580181181 | 0.406452516 | 0.4813 | 0.436418 |
| 100 | W | 0.352564017 | 0.545987665 | 0.479280744 | 0.4732 | 0.468506 |
| 101 | E | 0.312119384 | 0.488190352 | 0.543586861 | 0.4633 | 0.469511 |
| 102 | S | 0.331314146 | 0.522502544 | 0.578262734 | 0.4528 | 0.48891 |
| 103 | I | 0.341499119 | 0.548539342 | 0.638309363 | 0.4448 | 0.516684 |
| 104 | P | 0.338635361 | 0.573922764 | 0.69398788 | 0.4386 | 0.543737 |
| 105 | K | 0.336882784 | 0.594945001 | 0.776715771 | 0.4295 | 0.528964 |

| | | | | | | |
|---|---|---|---|---|---|---|
| 106 | K | 0.374143507 | 0.591041009 | 0.814458322 | 0.4231 | 0.540579 |
| 107 | F | 0.396585685 | 0.636506028 | 0.841296346 | 0.4146 | 0.534275 |
| 108 | K | 0.370475139 | 0.704863695 | 0.864736188 | 0.4019 | 0.482159 |
| 109 | P | 0.412738075 | 0.787054915 | 0.885640943 | 0.3881 | 0.443457 |
| 110 | L | 0.455037323 | 0.868895496 | 0.911610172 | 0.3722 | 0.424691 |
| 111 | S | 0.501287053 | 0.783451224 | 0.911611674 | 0.3587 | 0.373701 |
| 112 | N | 0.552090808 | 0.749562409 | 0.911613808 | 0.3453 | 0.320204 |
| 113 | R | 0.558111406 | 0.720774545 | 0.91254469 | 0.3353 | 0.293506 |
| 114 | I | 0.644930137 | 0.692486995 | 0.940719978 | 0.3243 | 0.263141 |
| 115 | N | 0.672129113 | 0.682111854 | 0.999999268 | 0.3101 | 0.226816 |
| 116 | V | 0.700023502 | 0.706707541 | 0.888888157 | 0.2948 | 0.22159 |
| 117 | I | 0.722572059 | 0.727799996 | 0.818504176 | 0.2793 | 0.218637 |
| 118 | L | 0.642633128 | 0.749370477 | 0.70739318 | 0.267 | 0.211027 |
| 119 | S | 0.552047034 | 0.771515017 | 0.596531908 | 0.2517 | 0.216564 |
| 120 | R | 0.449205164 | 0.856898493 | 0.485420797 | 0.233 | 0.264748 |
| 121 | T | 0.353540092 | 0.884731057 | 0.374310184 | 0.2158 | 0.321379 |
| 122 | L | 0.34779927 | 0.907569453 | 0.263199197 | 0.2002 | 0.385197 |
| 123 | K | 0.260149134 | 0.929882871 | 0.152088086 | 0.1901 | 0.468153 |
| 124 | K | 0.193320582 | 0.94915867 | 0.040976975 | 0.1839 | 0.532017 |
| 125 | E | 0.141520409 | 0.937835908 | 0.147171946 | 0.1799 | 0.523993 |
| 126 | D | 0.083890923 | 0.903341721 | 0.106444826 | 0.1744 | 0.485741 |
| 127 | F | 0.060199582 | 0.871311038 | 0.106444827 | 0.1696 | 0.438019 |
| 128 | D | 0.042268897 | 0.844691963 | 0.106194989 | 0.1697 | 0.364503 |
| 129 | E | 0.038224514 | 0.76994232 | 0.106194989 | 0.171 | 0.290967 |
| 130 | D | 0.031224162 | 0.685927299 | 0.106194989 | 0.1728 | 0.251408 |
| 131 | V | 0.029058384 | 0.600482788 | 0.106194989 | 0.1738 | 0.202758 |
| 132 | Y | 0.024949591 | 0.509895587 | 0.106194989 | 0.171 | 0.156901 |

| | | | | | | |
|---|---|---|---|---|---|---|
| 133 | I | 0.01926959 | 0.416209842 | 0.106194989 | 0.1684 | 0.133232 |
| 134 | I | 0.013779678 | 0.328083849 | 1.82E-08 | 0.1642 | 0.119891 |
| 135 | N | 0.009917082 | 0.25812849 | 7.80E-09 | 0.1584 | 0.100968 |
| 136 | K | 0.00853927 | 0.186391603 | 0 | 0.1547 | 0.096671 |
| 137 | V | 0.007431587 | 0.114206686 | 0 | 0.1474 | 0.098377 |
| 138 | E | 0.00632186 | 0.077945856 | 0 | 0.1448 | 0.08451 |
| 139 | D | 0.00544686 | 0.056923265 | 0 | 0.1427 | 0.073106 |
| 140 | L | 0.002938426 | 0.037206229 | 0 | 0.1394 | 0.069276 |
| 141 | I | 0.002373007 | 0.022656474 | 0 | 0.1387 | 0.062852 |
| 142 | V | 0.001855926 | 0.014057232 | 0 | 0.1375 | 0.057771 |
| 143 | L | 0.000243185 | 0.005323439 | 9.62E-05 | 0.1382 | 0.059961 |
| 144 | L | 2.83E-05 | 0.005192456 | 0.000115528 | 0.1368 | 0.057933 |
| 145 | G | 1.41E-05 | 0.00519244 | 0.000115605 | 0.1371 | 0.051636 |
| 146 | K | 1.11E-05 | 0 | 0.000124714 | 0.1382 | 0.048077 |
| 147 | L | 9.96E-06 | 2.60E-09 | 0.000124714 | 0.1368 | 0.043565 |
| 148 | N | 9.27E-06 | 2.60E-09 | 0.000124714 | 0.1351 | 0.039225 |
| 149 | Y | 0.000010197 | 2.70E-09 | 0.000124714 | 0.1334 | 0.040552 |
| 150 | Y | 1.01E-05 | 0.002258163 | 0.000124714 | 0.1292 | 0.042499 |
| 151 | K | 9.74E-06 | 0.03131853 | 0.000124714 | 0.125 | 0.042433 |
| 152 | C | 2.90E-05 | 0.138144421 | 2.85E-05 | 0.12 | 0.040686 |
| 153 | F | 4.70E-05 | 0.244286492 | 9.19E-06 | 0.1151 | 0.040933 |
| 154 | I | 6.42E-05 | 0.347790083 | 9.11E-06 | 0.1132 | 0.040382 |
| 155 | I | 7.53E-05 | 0.347791425 | 0 | 0.1134 | 0.041829 |
| 156 | G | 8.28E-05 | 0.347791423 | 0 | 0.1152 | 0.045695 |
| 157 | G | 9.48E-05 | 0.347792213 | 0 | 0.1152 | 0.051241 |
| 158 | S | 0.000105219 | 0.347792297 | 0 | 0.1154 | 0.058361 |
| 159 | V | 0.000120245 | 0.354837907 | 0.066990418 | 0.1136 | 0.070192 |

| | | | | | | |
|---|---|---|---|---|---|---|
| 160 | V | 0.000137627 | 0.325777561 | 0.066990418 | 0.1123 | 0.089168 |
| 161 | Y | 0.000134061 | 0.218951691 | 0.066990418 | 0.1088 | 0.115732 |
| 162 | Q | 0.000133707 | 0.11280963 | 0.066990418 | 0.1059 | 0.149732 |
| 163 | E | 0.00021765 | 0.009306042 | 0.066990418 | 0.1046 | 0.191339 |
| 164 | F | 0.000277985 | 0.009304756 | 0.071874979 | 0.1039 | 0.211661 |
| 165 | L | 0.000328778 | 0.009304797 | 0.073777986 | 0.1036 | 0.228118 |
| 166 | E | 0.00040118 | 0.00930409 | 0.083897688 | 0.1029 | 0.244656 |
| 167 | K | 0.000489413 | 0.00931206 | 0.194989685 | 0.104 | 0.252326 |
| 168 | K | 0.000574059 | 8.46E-06 | 0.239110089 | 0.1057 | 0.239903 |
| 169 | L | 0.001394407 | 1.60E-05 | 0.350202922 | 0.1042 | 0.223149 |
| 170 | I | 0.00142426 | 1.59E-05 | 0.350203605 | 0.1031 | 0.201522 |
| 171 | K | 0.002019769 | 1.59E-05 | 0.350203605 | 0.1032 | 0.183155 |
| 172 | K | 0.002637424 | 1.59E-05 | 0.350203605 | 0.1011 | 0.157258 |
| 173 | I | 0.002604525 | 0.000111324 | 0.345319044 | 0.099 | 0.129726 |
| 174 | Y | 0.002547701 | 0.001135743 | 0.343416036 | 0.0957 | 0.121653 |
| 175 | F | 0.002468075 | 0.001140957 | 0.333296334 | 0.095 | 0.112165 |
| 176 | T | 0.002462092 | 0.001138524 | 0.222204338 | 0.0961 | 0.109852 |
| 177 | R | 0.002456109 | 0.001161884 | 0.111093515 | 0.096 | 0.109736 |
| 178 | I | 0.001745793 | 0.001154579 | 6.82E-07 | 0.0998 | 0.111223 |
| 179 | N | 0.002227284 | 0.001155238 | 0 | 0.1038 | 0.114909 |
| 180 | S | 0.002360349 | 0.001155262 | 0 | 0.1073 | 0.121826 |
| 181 | T | 0.028520135 | 0.001913392 | 0 | 0.111 | 0.128442 |
| 182 | Y | 0.03951599 | 0.00181795 | 0 | 0.1134 | 0.120436 |
| 183 | E | 0.051246425 | 0.000793492 | 0 | 0.1167 | 0.117375 |
| 184 | C | 0.054335091 | 0.000788244 | 0 | 0.121 | 0.122108 |
| 185 | D | 0.057334285 | 0.00078263 | 0 | 0.1236 | 0.12295 |
| 186 | V | 0.061147987 | 0.00075911 | 0 | 0.1241 | 0.114864 |

| | | | | | | |
|---|---|---|---|---|---|---|
| 187 | F | 0.06355859 | 0.000758893 | 0 | 0.1223 | 0.13054 |
| 188 | F | 0.066240341 | 0.000758237 | 0 | 0.1228 | 0.161893 |
| 189 | P | 0.100657869 | 0.000759292 | 0 | 0.1215 | 0.175867 |
| 190 | E | 0.093345928 | 1.28E-06 | 2.80E-09 | 0.1214 | 0.202041 |
| 191 | I | 0.085551203 | 0.024512949 | 2.80E-09 | 0.1191 | 0.24011 |
| 192 | N | 0.078345275 | 0.107314328 | 2.80E-09 | 0.1188 | 0.261951 |
| 193 | E | 0.082300239 | 0.201637693 | 3.61E-06 | 0.1188 | 0.287546 |
| 194 | N | 0.101728099 | 0.250534856 | 3.61E-06 | 0.1199 | 0.333782 |
| 195 | E | 0.147339928 | 0.298849015 | 3.61E-06 | 0.1199 | 0.334662 |
| 196 | Y | 0.182749982 | 0.335344269 | 3.61E-06 | 0.1168 | 0.29873 |
| 197 | Q | 0.223338611 | 0.371694437 | 3.61E-06 | 0.1155 | 0.288221 |
| 198 | I | 0.231038406 | 0.434387382 | 3.61E-06 | 0.1125 | 0.271728 |
| 199 | I | 0.242177285 | 0.529309059 | 3.60E-06 | 0.1065 | 0.244583 |
| 200 | S | 0.242764183 | 0.598193318 | 3.60E-06 | 0.1031 | 0.24416 |
| 201 | V | 0.240491008 | 0.55640814 | 3.60E-06 | 0.0991 | 0.270312 |
| 202 | S | 0.241067559 | 0.501599356 | 0 | 0.0959 | 0.302876 |
| 203 | D | 0.236858375 | 0.454743846 | 0 | 0.094 | 0.311279 |
| 204 | V | 0.209603223 | 0.413745743 | 0 | 0.0931 | 0.34692 |
| 205 | Y | 0.178182091 | 0.377274295 | 0 | 0.0948 | 0.384289 |
| 206 | T | 0.155335455 | 0 | 0 | 0.0951 | 0.380604 |
| 207 | S | 0.129424453 | 0 | 0 | 0.0936 | 0.372956 |
| 208 | N | 0.110671681 | 0 | 0 | 0.094 | 0.353501 |
| 209 | N | 0.114620525 | 0 | 0 | 0.0922 | 0.289257 |
| 210 | T | 0.119467049 | 0 | 0 | 0.0899 | 0.228282 |
| 211 | T | 0.116522078 | 0 | 0 | 0.0874 | 0.213307 |
| 212 | L | 0.103613457 | 0 | 0 | 0.0848 | 0.204733 |
| 213 | D | 0.087806114 | 0 | 0 | 0.0838 | 0.230116 |

| | | | | | | |
|---|---|---|---|---|---|---|
| 214 | F | 0.083408773 | 0 | 0 | 0.0817 | 0.292583 |
| 215 | I | 0.063129218 | 0 | 0 | 0.0785 | 0.38098 |
| 216 | I | 0.048636683 | 0 | 0 | 0.0748 | 0.485324 |
| 217 | Y | 0.058120756 | 0 | 0 | 0.0712 | 0.591459 |
| 218 | K | 0.019176614 | 0 | 0 | 0.0678 | 0.641379 |
| 219 | K | 0.006051378 | 0 | 0 | 0.0617 | 0.677717 |
